# Long-term patterns of an interconnected core marine microbiota

**DOI:** 10.1101/2021.03.18.435965

**Authors:** Anders K. Krabberød, Ina M. Deutschmann, Marit F. M. Bjorbækmo, Vanessa Balagué, Caterina R. Giner, Isabel Ferrera, Esther Garcés, Ramon Massana, Josep M. Gasol, Ramiro Logares

## Abstract

**Background:** Ocean microbes constitute ∼70% of the marine biomass, are responsible for ∼50% of the Earth’s primary production, and are crucial for global biogeochemical cycles. Marine microbiotas include core taxa that are usually key for ecosystem function. Despite their importance, core marine microbes are relatively unknown, which reflects the lack of consensus on how to identify them. So far, most core microbiotas have been defined based on species occurrence and abundance. Yet, species interactions are also important to identify core microbes, as communities include interacting species. Here, we investigate interconnected bacteria and small protists of the core pelagic microbiota populating a long-term marine-coastal observatory in the Mediterranean Sea over a decade.

**Results:** Core microbes were defined as those present in >30% of the monthly samples over 10 years, with the strongest associations. The core microbiota included 259 Operational Taxonomic Units (OTUs) including 182 bacteria, 77 protists, and 1,411 strong and mostly positive (∼95%) associations. Core bacteria tended to be associated with other bacteria, while core protists tended to be associated with bacteria. The richness and abundance of core OTUs varied annually, decreasing in stratified warmers waters and increasing in colder mixed waters. Most core OTUs had a preference for one season, mostly winter, which featured subnetworks with the highest connectivity. Groups of highly associated taxa tended to include protists and bacteria with predominance in the same season, particularly winter. A group of 13 highly-connected hub-OTUs, with potentially important ecological roles dominated in winter and spring. Similarly, 18 connector OTUs with a low degree but high centrality were mostly associated with summer or autumn and may represent transitions between seasonal communities.

**Conclusions:** We found a relatively small and dynamic interconnected core microbiota in a model temperate marine-coastal site, with potential interactions being more deterministic in winter than in other seasons. These core microbes would be essential for the functioning of this ecosystem over the year. Other non-core taxa may also carry out important functions but would be redundant and non-essential. Our work contributes to the understanding of the dynamics and potential interactions of core microbes possibly sustaining ocean ecosystem function.

## BACKGROUND

Ecosystems are composed of interacting units embedded in and influenced by their physicochemical environment. Ecosystem function can be broadly defined as the biological, geochemical, and physical processes that occur within it. These processes will likely change or halt if specific organisms or gene-functions are removed, driving the ecosystem towards a new state or its collapse. It is hypothesized that ecological redundancy guarantees continuous ecosystem function, as multiple species could carry out the same or similar function [1]. And while the amount of functional redundancy in microbial ecosystems is a matter of debate [2, 3] it has also been observed that microbiotas in comparable habitats tend to share “core” species that are hypothesized to be fundamental for ecosystem function [4]. These core organisms and the functions they carry out might not be easily replaced.

Identifying the core microbiota is not straightforward as there are different ways of defining a core depending on the habitats and the questions being addressed [4]. One often-used approach is to identify species that tend to be recurrently present across spatiotemporal scales. This definition might not be sufficient, however, since communities are made up of interacting species [5]. A more appropriate definition of a core, therefore, needs to incorporate ecological interactions fundamental for the community in the location under study [4, 5]. This is particularly important in studies using DNA to investigate microbial communities, as a fraction of the detected taxa could be dormant, dead, or transient [6–8]. In the interaction-based definition taxa that do not appear to be interacting are excluded from the core [4].

Core microbiotas based on common presence have been widely studied in terrestrial animals, in particular humans [9] or cattle [10], as well in marine animals, in particular corals [11, 12] and sponges [13, 14]. Core microbiotas in non-host-associated systems, such as soils or the ocean, have been investigated to a lesser extent. In soils, for example, a global analysis identified a core group of 241 ubiquitous and dominant bacterial taxa with more or less invariant abundances and unclear habitat preferences [15]. In the tropical and subtropical global-ocean, a total of 68 bacteria and 57 picoeukaryotic operational taxonomic units (OTUs) have been identified that could be part of the core surface microbiota, as they were present in >80% of the globally-distributed samples [16].

Analyses of ocean time-series have also pointed to the existence of core microbiotas. For example, Gilbert et al. [17] investigated the microbiota of the English Channel for 6 years and found 12 abundant OTUs that were detected throughout the entire dataset (72 time-points), totaling ∼35% of the sequence abundance. Potentially core bacterial OTUs were detected in the SPOT time-series (southern California), in a study covering 10 years of monthly samples in the euphotic zone [18]. These potentially-core bacterial OTUs were present in >75% of the months, represented ∼7% (25-28 OTUs depending on depth) of the total richness, and had a high (>10%) relative abundance [18].

These studies have provided substantial insights on core marine microbiotas, although they typically define them in terms of species occurrence or abundance over spatiotemporal scales, rather than on potential interactions. As in other ecosystems, microbial interactions are essential for the functioning of the ocean ecosystem, where they guarantee the transfer of carbon and energy to upper trophic levels, as well as the recycling of carbon and nutrients [19]. Despite their importance, most microbial interactions in the ocean remain unknown [20]. A recent literature survey spanning the last 150 years indicated that we have documented a minor fraction of protist interactions in the ocean [21] and most likely, the same is true if not worse for bacteria.

During the last decade, association networks have been used to bridge this knowledge gap. Association networks are based on correlations between species’ abundances and they may reflect microbial interactions [22]. Contemporaneous positive correlations may point to interactions such as symbiosis, or similar niche preferences, while negative correlations may suggest predation, competition, or opposite niche preferences [23]. So far, network analyses have produced hypotheses on microbial interactions at the level of individual species across diverse ecosystems [22, 24, 25], a few of which have been experimentally validated [26]. In addition, networks can help detect species that have relatively more associations to other species (“hubs”), or species that connect different subgroups within a network, and which therefore may have important roles in the ecosystem. Groups of highly associated species in the network (“modules”) may represent niches [27, 28], and the amount of these modules may increase with increasing environmental selection [22]. Networks can also produce ecological insight at the community level, since their architecture can reflect community processes, such as selection [27].

Network analyses have been particularly useful for the investigation of microbial interactions in the ocean [25, 29]. A surface global-ocean network analysis of prokaryotes and single-celled eukaryotes indicated that ∼72% of the associations between microbes were positive and that most associations were between single-celled eukaryotes belonging to different organismal size-fractions [26]. Other studies using networks have indicated a limited number of associations between marine microbes and abiotic environmental variables [17, 18, 23, 26, 30–32], suggesting that microbial interactions have an important role in driving community turnover [32]. Despite the important insights these studies have provided, most of them share the limitation that they do not disentangle whether microbial associations may represent ecological interactions or environmental preferences [22].

Even though association networks based on long-term species dynamics may allow a more accurate delineation of core marine microbiotas, few studies have identified them in this manner. Consequently, we have a limited understanding of the interconnected set of organisms that may be key for ocean ecosystem function. Here we identify and investigate the core microbiota occurring in the marine-coastal Blanes Bay Microbial Observatory (Northwestern Mediterranean Sea) over 10 years. We delineated the core microbiota stringently, using potential interactions based on species abundances. We also made an effort to disentangle environmental effects in association networks by identifying and removing species associations that are a consequence of shared environmental preference and not interactions between the species [33]. We analyzed bacteria and protists from the pico- (0.2-3 µm) and nanoplankton (3-20 µm) organismal size fractions, which show a strong seasonality in this location [34–36]. Taxa relative abundances were estimated by sequencing the 16S and 18S rRNA-gene and delineating OTUs as Amplicon Sequence Variants (ASVs). Specifically, we ask: What taxa constitute the interconnected core microbiota and what are the main patterns of this assemblage over 10 years? Does the core microbiota feature seasonal sub-groups of highly associated species? What degree of association do bacteria and microbial eukaryotes have and do they show comparable connectivity? Can we identify core OTUs with central positions in the network that could have important ecological roles?

## RESULTS

### Composition and dynamics of the resident microbiota

Based on the data set containing 2,926 OTUs, (1,561 bacteria and 1,365 microbial eukaryotes) we first defined the resident OTUs as the bacteria and microbial eukaryotes present in >30% of the samples, which equals 36 out of 120 months (not necessarily consecutive). This threshold was selected as it includes seasonal OTUs that would be present recurrently in at least one season. The residents consisted of 709 OTUs: 354 Bacteria (∼54% relative read abundance) and 355 Eukaryotic OTUs (∼46% relative read abundance) [**Table 1**, see methods for calculation of relative read abundance]. The most abundant resident bacteria OTUs belonged to Oxyphotobacteria (mostly *Synechococcus*; ∼15% of total relative read abundance), Alphaproteobacteria (mostly SAR11 Clade Ia [∼9%, and clade II [∼4%]), and Gammaproteobacteria (mainly SAR86; ∼2%). The most abundant resident protist OTUs belonged to Dinophyceae (predominantly an unclassified dinoflagellate lineage [∼7%], Syndiniales Group I Clade 1 [∼7%] and *Gyrodinium* [∼4%]), Chlorophyta (mostly *Micromonas* [∼3%] and *Bathycoccus* [∼2%]), Ochrophyta (predominantly Mediophyceae [∼2%] and *Chaetoceros* [∼1%]) and Cryptophyceae (mainly a Cryptomonadales lineage [∼2%]) [**Figure 3, Table S1, Additional file 1**].

**Table 1.**
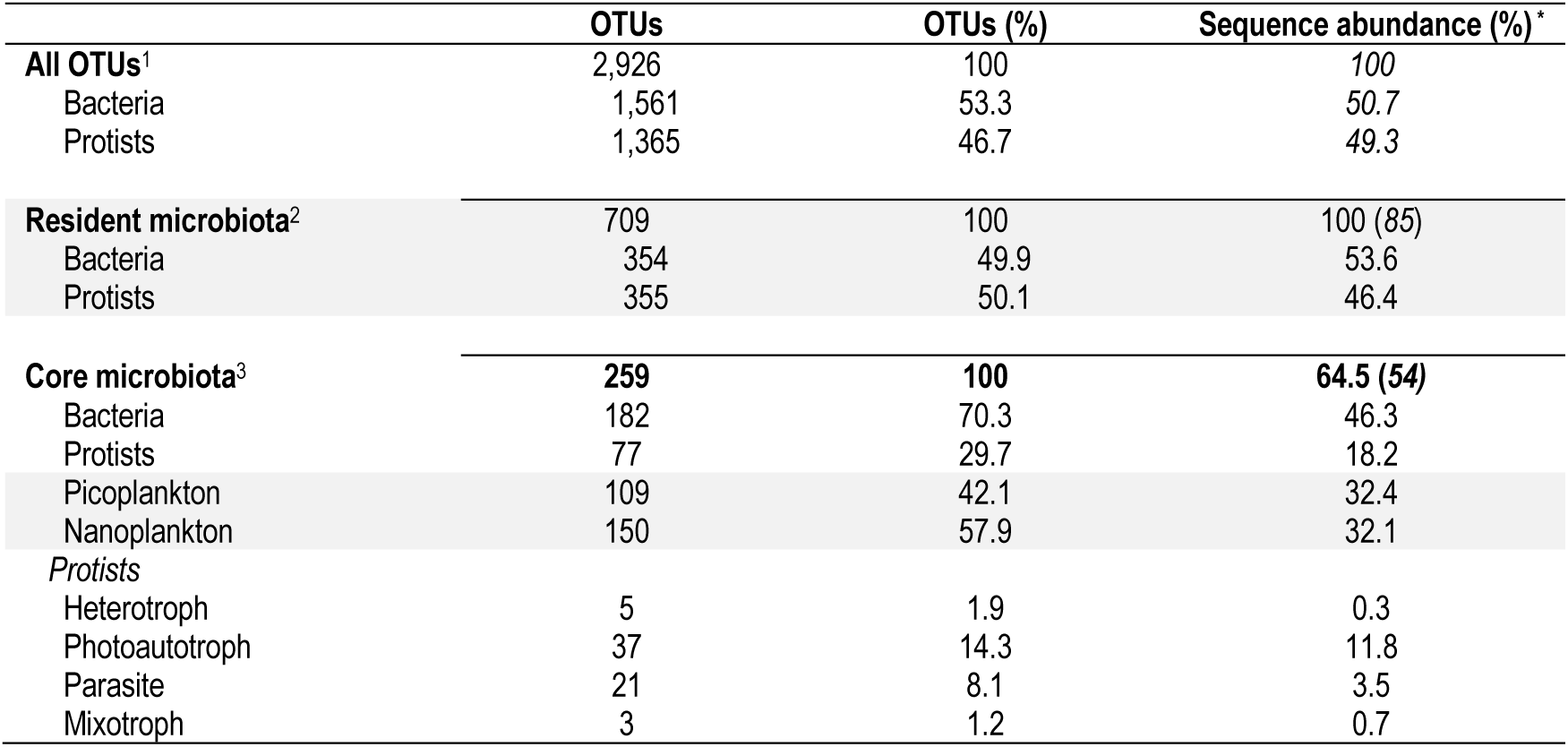
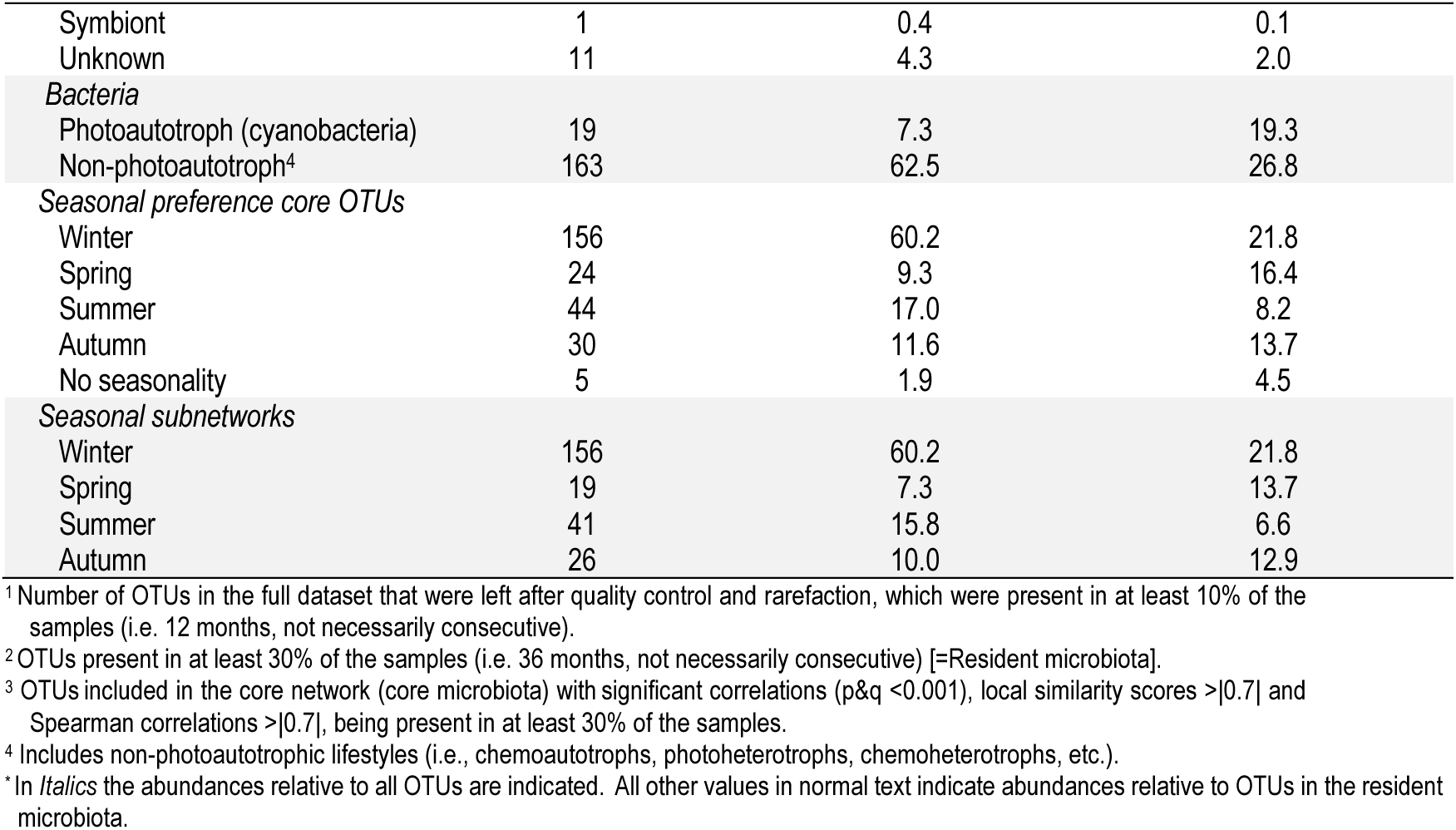
Description of the datasets.

|  | OTUs | OTUs (%) | Sequence abundance (%)<sup>*</sup> |
| --- | --- | --- | --- |
| **All OTUs<sup>1</sup>** | 2,926 | 100 | 100 |
| Bacteria | 1,561 | 53.3 | 50.7 |
| Protists | 1,365 | 46.7 | 49.3 |
| **Resident microbiota<sup>2</sup>** | 709 | 100 | 100 (85) |
| Bacteria | 354 | 49.9 | 53.6 |
| Protists | 355 | 50.1 | 46.4 |
| **Core microbiota<sup>3</sup>** | **259** | **100** | **64.5 (54)** |
| Bacteria | 182 | 70.3 | 46.3 |
| Protists | 77 | 29.7 | 18.2 |
| Picoplankton | 109 | 42.1 | 32.4 |
| Nanoplankton | 150 | 57.9 | 32.1 |
| *Protists* |  |  |  |
| Heterotroph | 5 | 1.9 | 0.3 |
| Photoautotroph | 37 | 14.3 | 11.8 |
| Parasite | 21 | 8.1 | 3.5 |
| Mixotroph | 3 | 1.2 | 0.7 |
| Symbiont | 1 | 0.4 | 0.1 |
| Unknown | 11 | 4.3 | 2.0 |
| *Bacteria* |  |  |  |
| Photoautotroph (cyanobacteria) | 19 | 7.3 | 19.3 |
| Non-photoautotroph<sup>4</sup> | 163 | 62.5 | 26.8 |
| *Seasonal preference core OTUs* |  |  |  |
| Winter | 156 | 60.2 | 21.8 |
| Spring | 24 | 9.3 | 16.4 |
| Summer | 44 | 17.0 | 8.2 |
| Autumn | 30 | 11.6 | 13.7 |
| No seasonality | 5 | 1.9 | 4.5 |
| *Seasonal subnetworks* |  |  |  |
| Winter | 156 | 60.2 | 21.8 |
| Spring | 19 | 7.3 | 13.7 |
| Summer | 41 | 15.8 | 6.6 |
| Autumn | 26 | 10.0 | 12.9 |

<sup>1</sup> Number of OTUs in the full dataset that were left after quality control and rarefaction, which were present in at least 10% of the samples (i.e. 12 months, not necessarily consecutive).
<sup>2</sup> OTUs present in at least 30% of the samples (i.e. 36 months, not necessarily consecutive) [=Resident microbiota].
<sup>3</sup> OTUs included in the core network (core microbiota) with significant correlations ( $p \leq 0.001$ ), local similarity scores $> 0.7$ and Spearman correlations $> 0.7$ , being present in at least 30% of the samples.
<sup>4</sup> Includes non-photoautotrophic lifestyles (i.e., chemoautotrophs, photoheterotrophs, chemoheterotrophs, etc.).
\* In *Italics* the abundances relative to all OTUs are indicated. All other values in normal text indicate abundances relative to OTUs in the resident microbiota.

The resident microbiota, including both protists and bacteria, showed seasonal variation over 10 years, with communities from the same season but different years tending to group (**Figure 1C and D**). The structure of the resident microbiota correlated to specific environmental variables during winter (nutrients, Total photosynthetic nanoflagellates [PNF; 2-5µm size], and small PNF [2µm]), spring (Total Chlorophyll a [Chla]), summer (daylength, temperature, Secchi disk depth and, the cell abundances of *Synechococcus,* Heterotrophic prokaryotes [HP] and Heterotrophic nanoflagellates [HNF, 2-5µm]) and autumn (salinity) [**Figure 1C**]. The environmental variables most relevant for explaining the variance of the resident microbiota were determined by stepwise model selection and distance-based redundancy analyses (dbRDA) [**Figure 1D**], leading to a dbRDA constrained and unconstrained variation of 41% and 59% respectively (**Figure 1D**). The selected variables were predominantly aligned with the axis summer (daylength, temperature, and the cell abundance of *Synechococcus* and HP) - winter (SiO_2_, small PNF [**Figure 1D**]. This dbRDA axis had the highest eigenvalue, explaining ∼55% of the constrained variation (**Figure 1D**). Even though the measured environmental variables did not explain the majority of the variation of the resident microbiota, they could account for a substantial fraction. This was further supported by Adonis analyses, which indicated that the measured environmental variables could explain ∼45% of the resident microbiota variance, with temperature and daylength having a predominant role by accounting for 30% of this variance (15% each).

**Figure 1.**
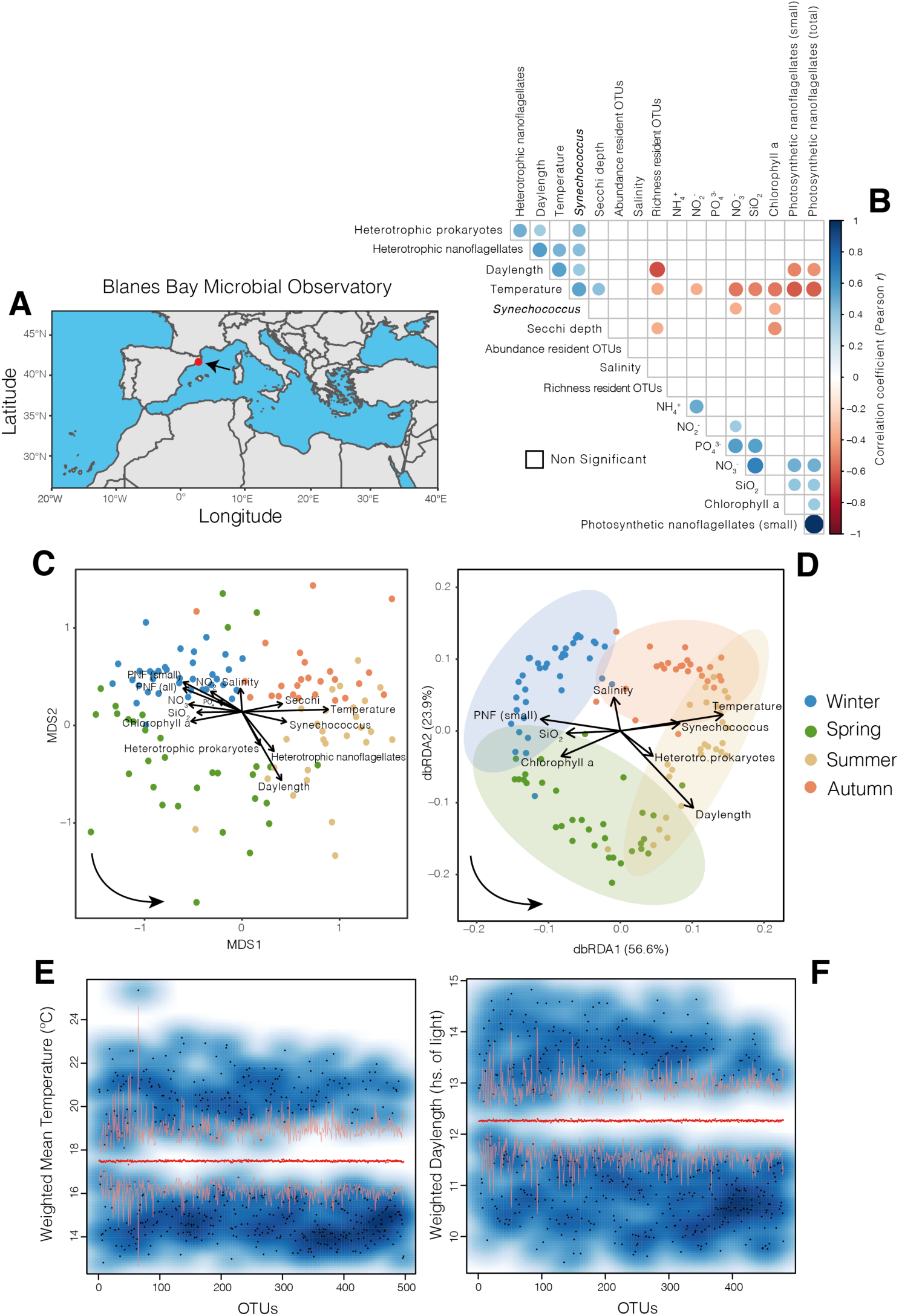
The Blanes Bay Microbial Observatory and the variation of its resident microbiota and measured environmental variables over ten years. **A)** Location of the Blanes Bay Microbial Observatory. **B)** All possible correlations between the measured environmental variables including the richness and abundance of resident OTUs (NB: only 709 resident OTUs are considered, see **Table 1**). Only significant Pearson correlation coefficients are shown (p<0.01). The p-values were corrected for multiple inference (Holm’s method). **C)** Unconstrained ordination (NMDS based on Bray Curtis dissimilarities) of communities including resident OTUs only, to which environmental variables were fitted. Only variables with a significant fit are shown (P<0.05). Arrows indicate the direction of the gradient and their length represents the strength of the correlation between resident OTUs and a particular environmental variable. The color of the samples (circles) indicates the season to which they belong. The bottom-left arrow indicates the direction of the seasonal change. PNF = photosynthetic nanoflagellates. **D)** Constrained ordination (Distance-based redundancy analyses, dbRDA, using Bray Curtis dissimilarities) including only the most relevant variables after stepwise model selection using permutation tests. Each axis (i.e., dbRDA1 and dbRDA2) indicates the amount of variance it explains according to the associated eigenvalues. The color of the samples (circles) indicates the season to which they belong. Arrows indicate the direction of the gradient and their length represents the strength of the correlation between resident OTUs and a particular environmental variable. The bottom-left arrow indicates the direction of the seasonal change. E-F) Resident OTUs displaying different niche preferences (blueish areas) in terms of the two most important abiotic variables: Temperature E) and Daylength F). The red dots indicate the randomization mean, and the orange curves represent the confidence limits. Black dots indicate individual OTUs for which temperature or daylength preferences are significantly (p<0.05) higher or lower than a random distribution over 10 years. At least two assemblages with different niches become evident: one preferring higher temperature and longer days (summer/spring), and another one preferring lower temperature and shorter days (winter/autumn). Note that several OTUs associated to Spring or Autumn are not expected to be detected with this approach, as their preferred temperature or daylength may not differ significantly from the randomized mean.

We then investigated whether temperature and daylength could determine the main niches. We found that ∼70% and ∼68% of the OTUs in the resident microbiota had niche preferences associated with temperature or daylength respectively (**Figure 1E-F**; Note that several OTUs preferring Spring or Autumn are not expected to be detected with this approach, as their preferred temperature or daylength may not differ significantly from the randomized mean). In total, 371 OTUs from the resident microbiota had both a temperature and a daylength niche preference that departed significantly from the randomization mean (**Figure 1E-F**). These 371 OTUs represented ∼52% of all OTUs in the resident microbiota, corresponding to ∼90% of the sequence abundance. In particular, 248 OTUs had a weighted mean for both temperature and daylength below the randomization mean (corresponding to winter/autumn), while 116 OTUs had a weighted mean above the randomization mean for both variables (corresponding to summer/spring). Interestingly, 7 OTUs displayed a weighted mean above and below the randomized mean for temperature and daylength respectively (corresponding to autumn or spring).

### Core network

To determine the core microbiota that incorporates possible interactions, we constructed an association network based on the resident OTUs and removed all OTUs that were not involved in strong and significant associations with any other OTUs. Specifically, we kept only the associations (edges in the network) with Local similarity score |LS| > 0.7, a false discovery rate adjusted p-value < 0.001 and Spearman |r| > 0.7. In addition, we removed all associations that seemed to be caused by environmental preferences of OTUs (see Methods). The core network consisted of 1,411 significant and strong correlations (**Figure 2A**) and was substantially smaller than the network based on the resident OTUs without stringent cut-offs (**Figure S1A, Additional file 2**, removed edges in **Figure S1B, Additional file 2**). The core network includes only the strongest microbial associations that are inferred during a decade and, according to our definition, determines the core microbiota. The associations in the core microbiota may represent proxies for species interactions since steps have been taken to remove associations that are driven by environmental factors.

**Figure 2.**
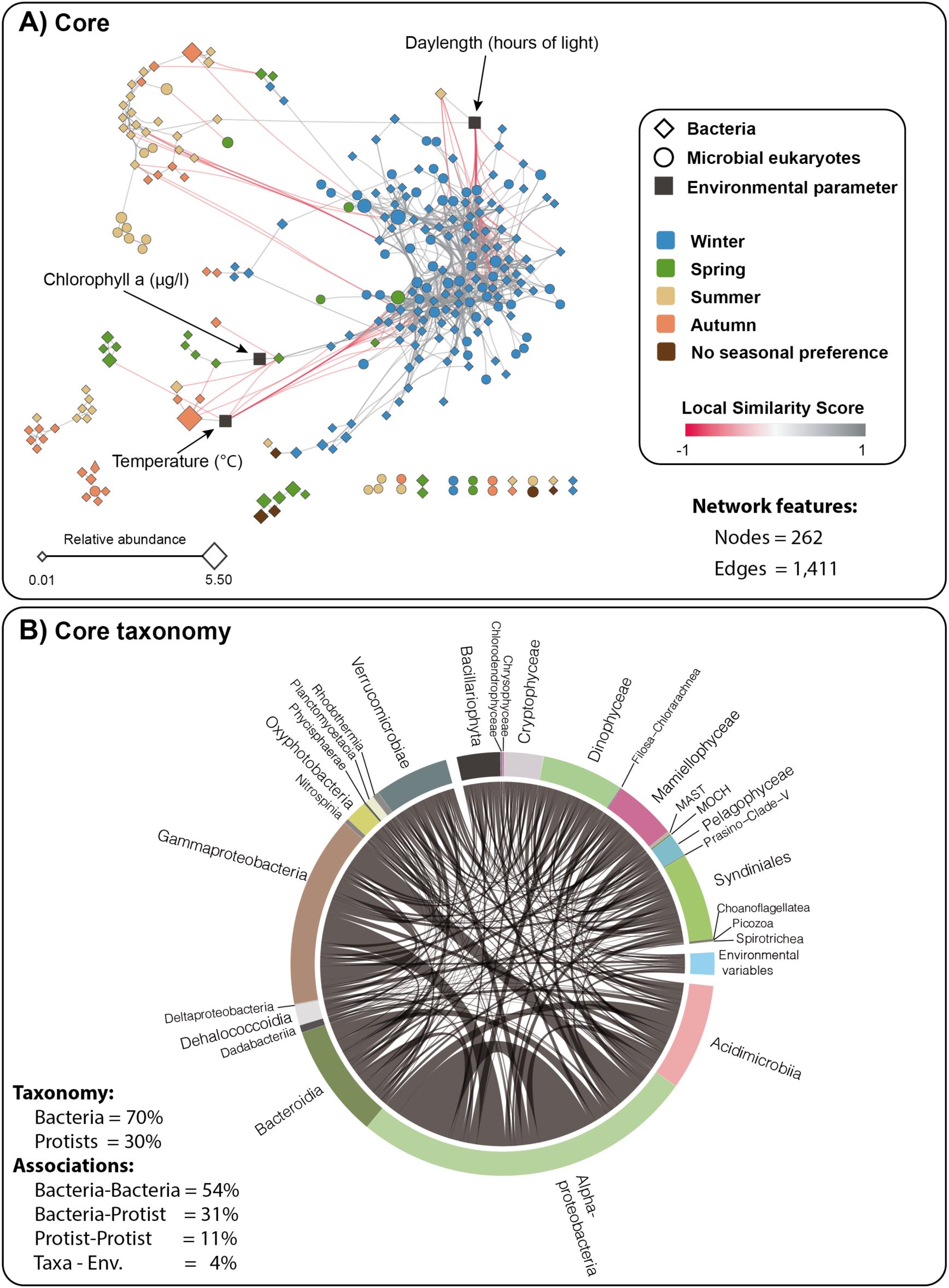
Core microbiota resulting from 10 years of monthly pico- and nanoplankton relative abundances. A) Core network including bacteria and microbial eukaryotic OTUs that occur ≥ 30% of the time during the studied decade (i.e. resident microbiota), with highly significant and strong associations (P<0.001 and Q<0.001, absolute local similarity score |LS| > 0.7, Spearman correlation |ρ|>0.7), where detected environmentally-driven edges were removed. The color of the edges (links) indicates whether the association is positive (grey) or negative (red). The shape of nodes indicates bacteria (rhomboid) or microbial eukaryotes (circle), and the color of nodes represents species seasonal preferences, determined using the indicator value (*indval*, p<0.05). Node size indicates OTU relative abundance. B) Core network as a Circos plot, indicating the high-rank taxonomy of the core OTUs. Since 95% of the associations are positive (see Table 2), we do not indicate whether an edge is positive or negative.

In the core network, most associations were positive (∼95%), pointing to the dominance of co-existence or symbiotic associations (**Table 2, Figure 2A**). The core network had “small world” properties [37], with a small average path length (i.e. number of nodes between any pair of nodes through the shortest path) and a relatively high clustering coefficient, showing that nodes tend to be connected to other nodes, forming tightly knit groups, more than what it would be expected by chance (**Table 3**). Since node degree was not correlated with OTU abundance (**Figure S2, Additional file 3**), the associations between OTUs are not caused by a high sequence abundance alone, as the most abundant OTUs did not tend to be the most connected.

**Table 2.** Core associations. See **Figure 2**.

|  | Association #<br>(edges) | Co-occurrences<br>(positive) | Co-exclusions<br>(negative) |
| --- | --- | --- | --- |
| All | 1,411 | 1,341 (95.0%) | 70 (5.0%) |
| Within Picoplankton | 378 | 353 (93.3%) | 25 (6.6%) |
| Within Nanoplankton | 791 | 748 (94.6%) | 43 (5.4%) |
| Picoplankton-Nanoplankton | 242 | 240 (99.2%) | 2 (0.8%) |

**Table 3.**
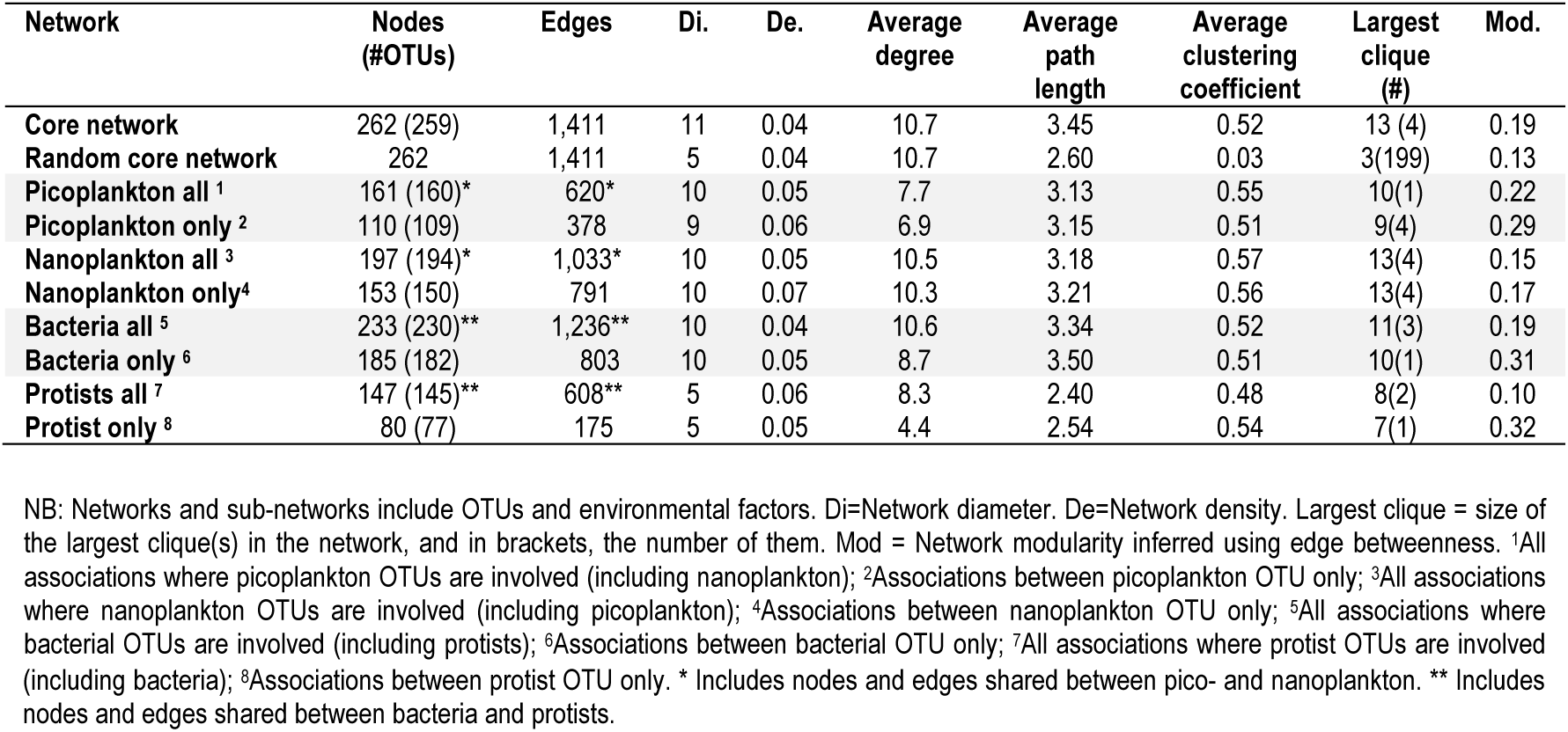
Core network and sub-networks statistics.

| Network | Nodes<br>(#OTUs) | Edges | Di. | De. | Average<br>degree | Average<br>path<br>length | Average<br>clustering<br>coefficient | Largest<br>clique<br>(#) | Mod. |
| --- | --- | --- | --- | --- | --- | --- | --- | --- | --- |
| Core network | 262 (259) | 1,411 | 11 | 0.04 | 10.7 | 3.45 | 0.52 | 13 (4) | 0.19 |
| Random core network | 262 | 1,411 | 5 | 0.04 | 10.7 | 2.60 | 0.03 | 3(199) | 0.13 |
| Picoplankton all <sup>1</sup> | 161 (160)* | 620* | 10 | 0.05 | 7.7 | 3.13 | 0.55 | 10(1) | 0.22 |
| Picoplankton only <sup>2</sup> | 110 (109) | 378 | 9 | 0.06 | 6.9 | 3.15 | 0.51 | 9(4) | 0.29 |
| Nanoplankton all <sup>3</sup> | 197 (194)* | 1,033* | 10 | 0.05 | 10.5 | 3.18 | 0.57 | 13(4) | 0.15 |
| Nanoplankton only <sup>4</sup> | 153 (150) | 791 | 10 | 0.07 | 10.3 | 3.21 | 0.56 | 13(4) | 0.17 |
| Bacteria all <sup>5</sup> | 233 (230)** | 1,236** | 10 | 0.04 | 10.6 | 3.34 | 0.52 | 11(3) | 0.19 |
| Bacteria only <sup>6</sup> | 185 (182) | 803 | 10 | 0.05 | 8.7 | 3.50 | 0.51 | 10(1) | 0.31 |
| Protists all <sup>7</sup> | 147 (145)** | 608** | 5 | 0.06 | 8.3 | 2.40 | 0.48 | 8(2) | 0.10 |
| Protist only <sup>8</sup> | 80 (77) | 175 | 5 | 0.05 | 4.4 | 2.54 | 0.54 | 7(1) | 0.32 |

NB: Networks and sub-networks include OTUs and environmental factors. Di=Network diameter. De=Network density. Largest clique = size of the largest clique(s) in the network, and in brackets, the number of them. Mod = Network modularity inferred using edge betweenness. <sup>1</sup>All associations where picoplankton OTUs are involved (including nanoplankton); <sup>2</sup>Associations between picoplankton OTU only; <sup>3</sup>All associations where nanoplankton OTUs are involved (including picoplankton); <sup>4</sup>Associations between nanoplankton OTU only; <sup>5</sup>All associations where bacterial OTUs are involved (including protists); <sup>6</sup>Associations between bacterial OTU only; <sup>7</sup>All associations where protist OTUs are involved
(including bacteria); \*Associations between protist OTU only. \* Includes nodes and edges shared between pico- and nanoplankton. \*\* Includes nodes and edges shared between bacteria and protists.

The core network displayed a winter cluster, while no clear clusters could be defined for the other seasons (**Figure 2A**). Of the 15 environmental variables analyzed, only 3 were found to be significantly correlated with core OTUs: *daylength*, showing strong correlations with 33 OTUs, *temperature*, correlated with 14 OTUs, and *Chlorophyll a*, correlated with 1 OTU (**Figure 2A**). Therefore, the analysis of the core network also points to the importance of temperature and daylength in the decade-long seasonal dynamics of the studied microbial ecosystem. It is also coherent with the Adonis and ordination analyses (**Figure 1C-B**). However, the associations between these environmental parameters with taxa represented only 4% of all the associations (**Figure 2B**).

Of the 709 OTUs from the resident microbiota (**Figure 3**), only 259 OTUs (35%) were left in the core network (182 bacteria (∼70%) and 77 microbial eukaryotic OTUs (∼30%); **Table 1, Figure 2**). The monthly taxonomic composition of the resident microbiota differed from that of the core (**Figure 3**). The core OTUs accounted for ∼64% of the relative read abundance of the resident microbiota (**Table 1**). The core OTUs had annual variation in terms of richness and abundance over the 10 years for both the pico- and nanoplankton, with microbial eukaryotes decreasing markedly in OTU richness and relative read abundance in the warmer seasons, and increasing during colder periods (**Figure 3**).

**Figure 3.**
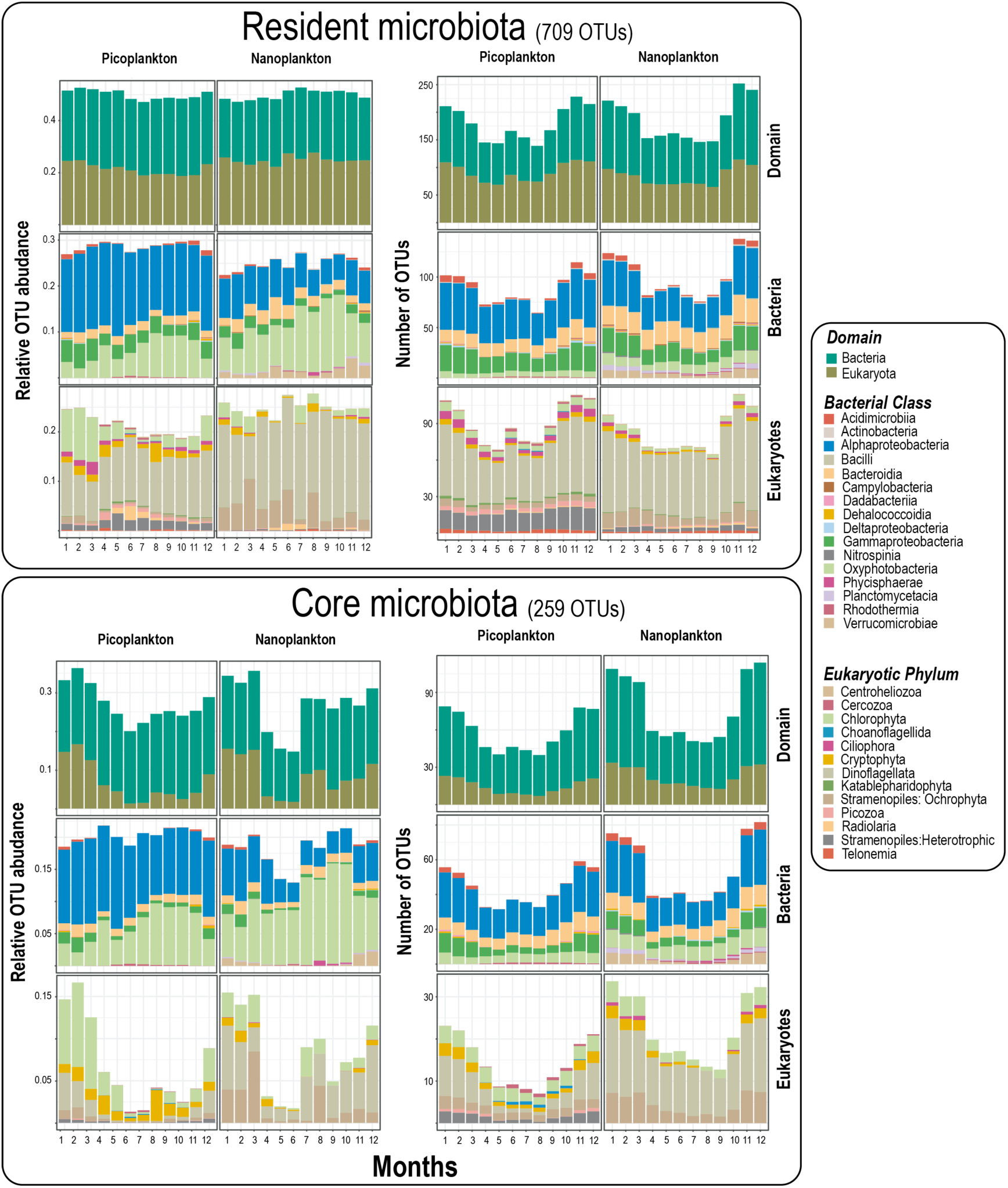
The monthly variation in the resident and core microbiotas over 10 years. *Upper panels:* The resident microbiota is defined as those eukaryotes and bacteria that occur in at least 30% of the samples over 10 years. The relative OTU abundance (left panel) and number of OTUs (right panel) for different domains and taxonomic levels in the resident microbiota are shown. Note that the relative abundance of Bacteria vs. Eukaryotes does not necessarily reflect organismal abundances on the sampling site, but the amplicon relative abundance after PCR. Relative abundances were calculated for each year and aggregated over the corresponding months along the 10 years for the resident microbiota, then split into size fractions (NB: relative abundance for both domains and size fraction sums up to 1 for each month across ten years). *Lower panels:* Core microbiota over 10 years. The relative abundances of core OTUs reflect the remaining proportions after removing all the OTUs that were not strongly associated when building networks. Relative OTU abundance (left panel) and number of OTUs (right panel) for different domains and taxonomic levels among the core OTUs.

The most abundant bacteria (**Figure 3**; **Table S2, Additional file 1**) among the core OTUs were Oxyphotobacteria (mostly *Synechococcus*), total abundance ∼14% of the resident microbiota, followed by Alphaproteobacteria, with SAR11 clades Ia and II representing ∼9% and ∼2% respectively. The most abundant microbial eukaryotic groups were *Micromonas*, *Bathycoccus*, Dinophyceae, and Cryptomonadales (each ∼2%) [**Figure 3**; **Table S3, Additional file 1**]. In terms of diversity and abundance, bacterial non-phototrophs (including chemoautotrophs, photoheterotrophs, chemoheterotrophs) were the most prevalent in the core microbiota, representing ∼62% of the OTUs and a quarter of the total relative read abundance (**Table 1**). In turn, protistan heterotrophs represented a minor fraction of the diversity and relative abundance **(Table 1)**. Bacteria photoautotrophs were relatively more abundant than their protistan counterparts but less diverse (**Table 1**). Protistan parasites represented ∼8% of the OTUs and ∼3% of the abundance, while the remaining protistan lifestyles had a minor relevance in the core microbiota (**Table 1**).

### Intra- and cross-domain core associations

Bacteria tended to be associated with other bacteria (**Table 3 & 4**; **Figure 2B**), with Bacteria-Bacteria associations making up ∼54% of all associations, while Protist-Protist associations accounted for 11% (**Table 4**). The connectivity of the bacterial subnetworks was higher (mean degree ∼10) than the protist counterparts (mean degree ∼6), regardless of whether these networks included exclusively bacteria, protists, or both (**Table 3**).

**Table 4.** Core associations within and between taxonomic domains and size fractions.

| Network | Association type<sup>1</sup> | # Associations |
| --- | --- | --- |
| **Core network** | Total | 1,411 |
|  | Bacteria - Bacteria | 767 (54%) |
|  | Bacteria - Protist | 433 (31%) |
|  | Protist - Protist | 161 (11%) |
|  | Environmental factor - Bacteria | 36 (3%) |
|  | Environmental factor - Protist | 14 (1%) |
| **Picoplankton subnetwork** | Total | 378 |
|  | Bacteria - Bacteria | 241 (64%) |
|  | Bacteria - Protist | 94 (25%) |
|  | Protist - Protist | 31 (8%) |
|  | Environmental factor - Bacteria | 12 (3%) |
|  | Environmental factor - Protist | 0 (0%) |
| **Nanoplankton subnetwork** | Total | 791 |
|  | Bacteria - Bacteria | 394 (50%) |
|  | Bacteria - Protist | 246 (31%) |
|  | Protist - Protist | 113 (14%) |
|  | Environmental factor - Bacteria | 24 (3%) |
|  | Environmental factor - Protist | 14 (2%) |

<sup>1</sup>"Bacteria – Bacteria" indicates associations between two bacterial OTUs. "Protist – Protist" are associations between two unicellular eukaryotes and "Bacteria – Protist" are associations between one eukaryote and one bacterial OTU. "Environmental factor – Protist" and "Environmental factor – Bacteria" are associations between an environmental factor and a eukaryotic or bacterial OTU.

In particular, there was a substantial number of associations between Alpha- and Gammaproteobacteria, between Alphaproteobacteria and Acidiimicrobia as well as among Alphaproteobacteria OTUs (**Figure 2B**). Eukaryotic OTUs did not show a similar trend with associations between OTUs of the same taxonomic ranks (**Figure 2B**). In terms of cross-domain associations, Alphaproteobacteria OTUs had several associations with most major protistan groups (i.e. dinoflagellates, diatoms, cryptophytes, Mamiellophyceae, and Syndiniales) [**Figure 2B**].

### Core associations within the pico- and within the nanoplankton

While the pico- and nano-size fractions indicate different lifestyles in bacteria (free-living or particle-attached), they indicate different cell sizes in protists, and this could be reflected in association networks. Nanoplankton sub-networks were larger and more connected than picoplankton counterparts (**Figure 4, Table 3**). This pattern was observed in both sub-networks considering associations from the same or both size fractions (**Table 3**). Nanoplankton sub-networks had a higher average degree (∼10) than picoplankton sub-networks (∼7; Wilcoxon p<0.05), while not differing much in other network statistics (**Table 3**). Most associations in the pico- and nanoplankton were positive (>93%), while the associations between OTUs from different size fractions represented only ∼17% of the total, being ∼99% positive (**Table 2**).

**Figure 4.**
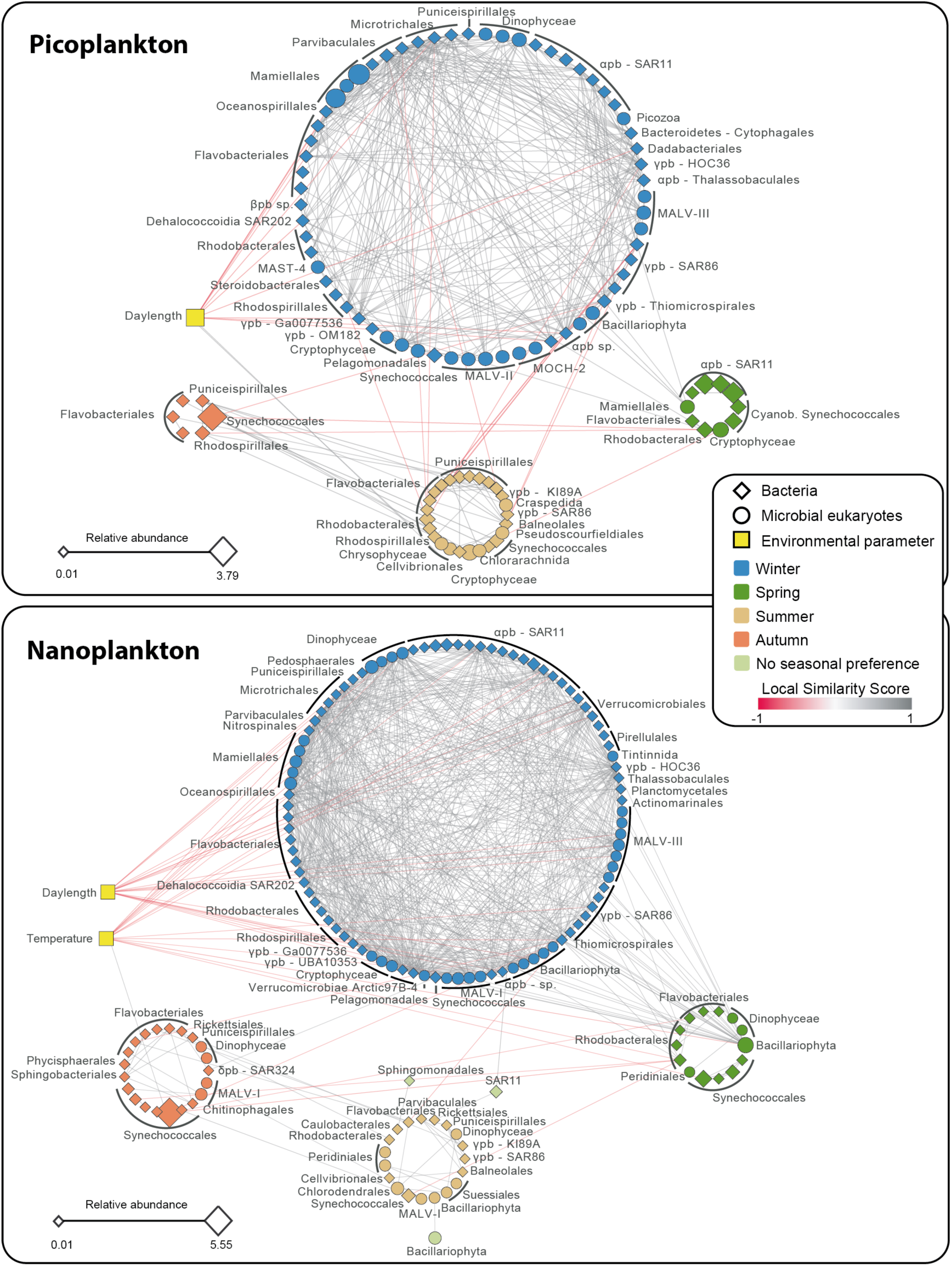
Pico- and nanoplankton core sub-networks. The shape of the nodes indicates bacteria (rhomboid) or microbial eukaryotes (circle), and the color of nodes represents species seasonal preferences, determined using the indicator value (p<0.05). The color of the edges indicates if the association is positive (grey) or negative (red). Node size indicates OTU relative abundance from the core microbiota.

In the pico- or nanoplankton sub-networks that include OTUs from the same size fraction, the number of bacterial core OTUs was higher than the protistan counterparts (103 bacterial vs. 47 protistan OTUs in the nanoplankton, and 79 bacterial vs. 30 protistan OTUs in the picoplankton) (**Figure 4, Table 3**). Still, core OTUs in both the pico- and nanoplankton had comparable sequence abundances: ∼27% of the resident microbiota in each size fraction. Within the picoplankton, 64% of the associations were between bacteria, 8% between eukaryotes, and 25% between eukaryotes and bacteria (**Table 4**). In turn, in the nanoplankton, 50% of the edges were between bacteria, 14% between eukaryotes, and 31% between eukaryotes and bacteria (**Table 4**). Overall, the BBMO pico- and nanoplankton sub-networks differed in size, connectivity, and taxonomic composition, while they were similar in terms of positive connections and relative sequence abundance.

### Network seasonality

The indicator value (IndVal) was used to infer the seasonal preference of core OTUs. Most of the core OTUs (98%; 254 out of 259 OTUs) showed a clear preference for one of the four seasons, pointing to a marked seasonality in the core microbiota (**Figure 4; Table 5; Tables S4 & S5, Additional file 1**). Winter had the highest quantity of core OTUs and the highest network connectivity (average degree ∼13), compared to the other seasons (average degrees ∼2 – ∼6) [**Figure 4**; **Table 5**]. The average path length was larger in the core network compared to a random network of the same size (**Table 3**). Yet, all sub-networks associated with size fractions and seasons (**Table 5**) had shorter path lengths than the random network, indicating that nodes tended to be connected within seasons and size fractions. This was also supported by an increase in network density when comparing the core network (**Table 3**) and the core network subdivided into seasons (**Table 5**), against the core network subdivided into both seasons and size fractions (**Table 5**). The five OTUs that did not show any seasonal preference, among them SAR11 Clades Ia & II, showed high to moderate abundances but had a low number of associations to other OTUs (**Tables S4, S5, S6, Additional file 1**). Thus, network connectivity in the BBMO appears to be heterogeneous over time, peaking in winter and remaining low in the other seasons.

**Table 5:** Subnetworks including core OTUs displaying seasonal preference.

|  | Sub-network | Number of OTUs | Edges | Di. | De. | Average degree | Average path length | Average clustering coefficient | Largest clique (#) | Mod. |
| --- | --- | --- | --- | --- | --- | --- | --- | --- | --- | --- |
| All | Winter | 156 | 1,175 | 7 | 0.10 | 15.1 | 2.62 | 0.54 | 13(4) | 0.19 |
|  | Spring | 19 | 16 | 4 | 0.09 | 1.7 | 1.56 | 0.44 | 4(1) | 0.75 |
|  | Summer | 41 | 56 | 7 | 0.07 | 2.7 | 2.90 | 0.49 | 6(1) | 0.53 |
|  | Autumn | 26 | 25 | 3 | 0.08 | 1.9 | 1.59 | 0.46 | 4(2) | 0.73 |
| Pico | Winter | 63 | 286 | 6 | 0.15 | 9.1 | 2.35 | 0.53 | 9(4) | 0.10 |
|  | Spring | 8 | 5 | 3 | 0.18 | 1.2 | 1.50 | 0.00 | 2(5) | 0.56 |
|  | Summer | 25 | 36 | 5 | 0.12 | 2.9 | 2.20 | 0.41 | 6(1) | 0.23 |
|  | Autumn | 5 | 3 | 2 | 0.30 | 1.2 | 1.25 | 0.00 | 2(3) | 0.44 |
| Nano | Winter | 92 | 658 | 6 | 0.16 | 14.3 | 2.40 | 0.61 | 13(4) | 0.04 |
|  | Spring | 11 | 11 | 4 | 0.20 | 2.0 | 1.59 | 0.57 | 4(1) | 0.56 |
|  | Summer | 13 | 17 | 3 | 0.22 | 2.6 | 1.70 | 0.65 | 4(1) | 0.50 |
|  | Autumn | 17 | 18 | 3 | 0.13 | 2.1 | 1.35 | 0.56 | 4(2) | 0.60 |

NB: Subnetworks include OTUs only. Di=Network diameter. De=Network density. Largest clique = size of the largest clique(s) in the network, and in brackets, the number of them. Mod = Network modularity inferred using edge betweenness.

### Groups of highly associated OTUs

Within the core network, we identified groups that were more connected to each other than to the rest of the network (called modules). These groups of OTUs may indicate recurring associations that are likely important for the stability of ecosystem function. We identified 12 modules in both the pico- and nanoplankton subnetworks (**Table S7, Additional file 1**). Modules tended to include OTUs from the same season (**Table S8, Additional file 1**), with main modules (i.e. MCODE score >4) including OTUs predominantly associated with winter, summer, and autumn (**Figure 5**). Overall, winter modules prevailed (5 out of 7) among the main modules (**Figure 5**), while modules with scores ≤ 4 did not tend to be associated with a specific season (**Table S8, Additional file 1**). Two main winter modules had members that were negatively correlated to temperature and daylength (**Figure 5**; Modules 1 and 4, nanoplankton).

**Figure 5.**
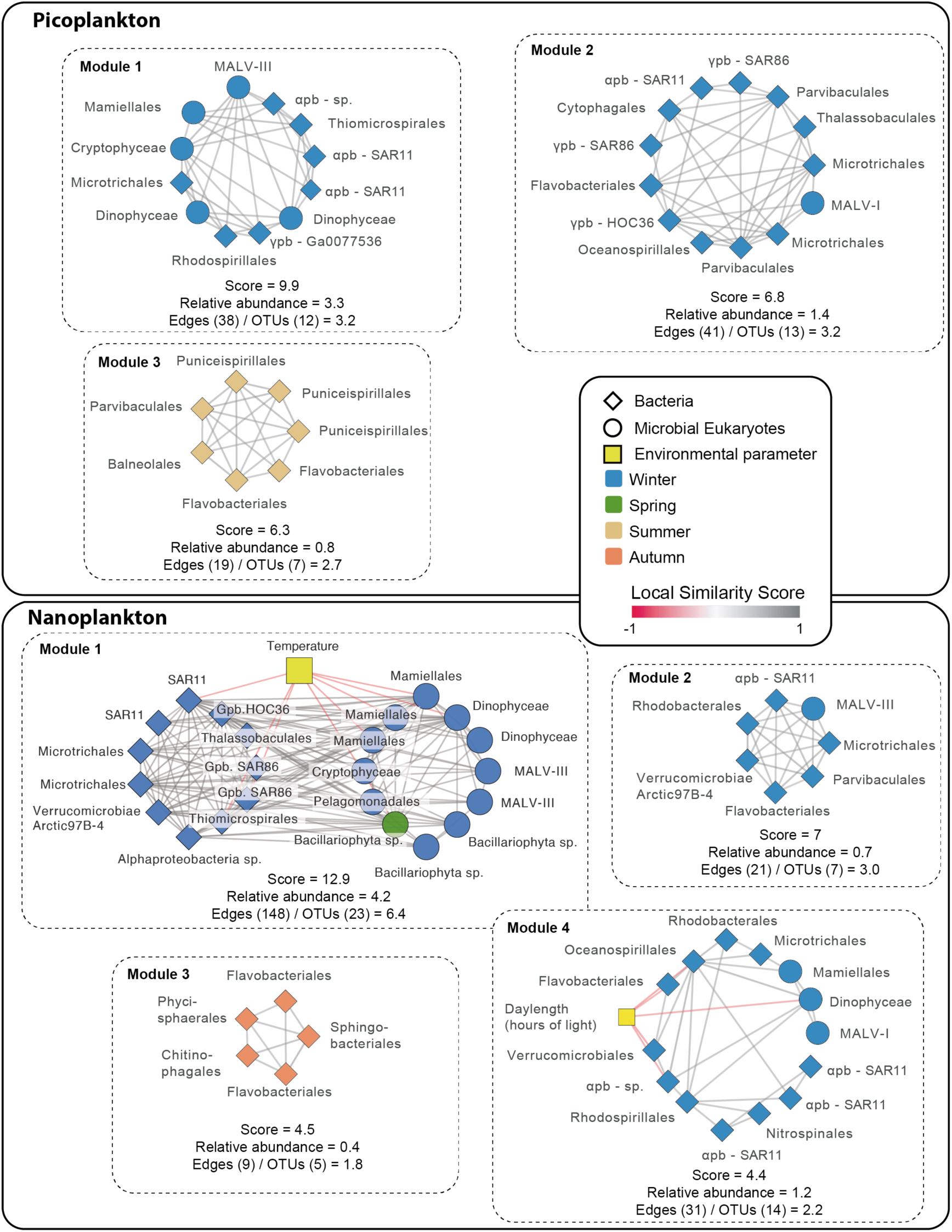
Main modules in the core network. Modules with MCODE score >4 are shown for picoplankton (upper panel) and nanoplankton (lower panel). For each module, the MCODE score and relative amplicon abundance of the taxa included in it (as % of the resident microbiota) are indicated. In addition, the numbers of edges and OTUs within the modules are shown as edges/OTUs; this quotient estimates the average number of edges per OTU within the different modules. The edges represent correlations with |LS| > 0.7, |ρ|>0.7, P<0.001 and Q<0.001. The color of the edges indicates positive (grey) or negative (red) associations. The shape of nodes indicates bacteria (rhomboid) or microbial eukaryotes (circle), and the color of nodes represents species seasonal preferences, determined using the indicator value (p<0.05). pb = Proteobacteria

The total relative sequence abundance of core OTUs included in modules was ∼24% (proportional to the resident microbiota), while the total abundance of individual modules ranged between ∼6% and ∼0.3% (**Table S7, Additional file 1**). In turn, the relative abundance of core OTUs included in modules ranged between 0.01% and ∼2% (**Table S8, Additional file 1**). In most modules, a few OTUs tended to dominate the abundance, although there were exceptions, such as module 4 of the picoplankton, where all SAR11 members featured abundances >1% (**Table S8, Additional file 1**). In addition, several OTUs within modules had relatively low abundances (**Table S8, Additional file 1**), supporting modules as a real feature of the network and not just the agglomeration of abundant taxa.

### Central OTUs

Biological networks typically contain nodes (i.e. OTUs) that hold more “central” positions in the network than others [22]. Even though the ecological role of these hub and connector OTUs is unclear, it is acknowledged that they could reflect taxa with important ecological functions [22]. There is no universal definition for hub or connector OTUs, yet, in this work, we have used stringent thresholds to determine them *ad hoc* (see Methods). We have identified 13 hub-OTUs that were associated with winter or spring (**Table 6**). Hubs did not include highly abundant OTUs, such as *Synechococcus* or SAR11 (**Table 6**), but instead, they included several OTUs with moderate-low abundance (<1%) and high degree (ranging between 26-60) [**Table 6**]. For example, the Gammaproteobacteria OTU bn_000226 had a relative abundance of 0.04% and a degree of 60 **(Table 6)**. Hubs included other moderately abundant OTUs, such as the eukaryotic picoalgae *Bathycoccus,* which was abundant in winter, as well as an unidentified dinoflagellate (**Table 6**).

**Table 6.** Central OTUs.

| OTU | Class | Lowest rank taxonomy | Relative Abundance (%)<sup>1</sup> | Degree | Betweenness Centrality | Closeness Centrality | Season |
| --- | --- | --- | --- | --- | --- | --- | --- |
| **Hubs** |  |  |  |  |  |  |  |
| en_00092 | Mamiellophyceae | *Bathycoccus* | 0.51 | 42 | 0.04 | 0.42 | Winter |
| en_00119 | Dinophyceae | - | 0.41 | 50 | 0.03 | 0.42 | Winter |
| bp_000037 | Alphaproteobacteria | Parvibaculales_OCS116 | 0.31 | 45 | 0.08 | 0.43 | Winter |
| bp_000039 | Gammaproteobacteria | SUP05_cluster | 0.28 | 29 | 0.12 | 0.41 | Spring |
| bn_000039 | Gammaproteobacteria | SUP05_cluster | 0.21 | 42 | 0.17 | 0.44 | Spring |
| bn_000037 | Alphaproteobacteria | Parvibaculales_OCS116 | 0.20 | 40 | 0.05 | 0.42 | Spring |
| bp_000059 | Gammaproteobacteria | SAR86 | 0.20 | 24 | 0.09 | 0.40 | Spring |
| ep_00070 | Cryptophyceae | Cryptomonadales_X | 0.13 | 40 | 0.04 | 0.42 | Winter |
| bn_000059 | Gammaproteobacteria | SAR86 | 0.12 | 24 | 0.03 | 0.40 | Spring |
| bn_000102 | Alphaproteobacteria | Nisaeaceae_OM75 | 0.09 | 26 | 0.03 | 0.38 | Winter |
| bp_000193 | Alphaproteobacteria | - | 0.06 | 37 | 0.03 | 0.40 | Winter |
| bn_000170 | Acidimicrobiia | Sva0996_marine_group | 0.06 | 59 | 0.06 | 0.44 | Winter |
| bn_000226 | Gammaproteobacteria | HOC36 | 0.04 | 60 | 0.06 | 0.43 | Winter |
| **Connectors** |  |  |  |  |  |  |  |
| bp_000001 | Oxyphotobacteria | *Synechococcus* (CC9902) | 3.79 | 5 | 0.05 | 0.30 | Autumn |
| bp_000002 | Alphaproteobacteria | SAR11 Clade_la | 2.26 | 2 | 0.40 | 0.56 | Spring |
| bp_000004 | Alphaproteobacteria | SAR11 Clade_la | 2.02 | 3 | 0.15 | 0.63 | NA |
| bp_000007 | Alphaproteobacteria | SAR11 Clade_la | 1.38 | 3 | 0.60 | 0.71 | NA |
| bp_000008 | Alphaproteobacteria | SAR11 Clade_la | 1.15 | 3 | 0.15 | 0.63 | NA |
| bn_000008 | Alphaproteobacteria | SAR11 Clade_la | 0.68 | 5 | 0.03 | 0.27 | Winter |
| en_00059 | Chlorodendrophyceae | *Tetraselmis* | 0.66 | 4 | 0.05 | 0.26 | Summer |
| bn_000020 | Oxyphotobacteria | - | 0.56 | 3 | 0.60 | 0.67 | Autumn |
| en_00161 | Syndiniales | Syndiniales-Group-I-Clade-4_X | 0.42 | 4 | 0.80 | 0.75 | Autumn |
| bn_000018 | Oxyphotobacteria | *Prochlorococcus* MIT9313 | 0.41 | 5 | 0.04 | 0.24 | Winter |
| bn_000054 | Alphaproteobacteria | Puniceispirillales_SAR116 | 0.11 | 4 | 0.14 | 0.40 | Autumn |
| bn_000062 | Alphaproteobacteria | Puniceispirillales_SAR116 | 0.08 | 3 | 0.55 | 0.50 | Autumn |
| bn_000077 | Rhodothermia | *Balneola* | 0.07 | 3 | 0.17 | 0.32 | Summer |
| bn_000112 | Gammaproteobacteria | KI89A | 0.06 | 4 | 0.53 | 0.48 | Summer |
| bn_000156 | Alphaproteobacteria | Parvibaculales_PS1 | 0.05 | 4 | 0.14 | 0.40 | Summer |
| bn_000281 | Bacteroidia | Sphingobacteriales_NS11-12 | 0.05 | 5 | 0.16 | 0.44 | Autumn |
| bn_000221 | Alphaproteobacteria | Puniceispirillales_SAR116 | 0.04 | 5 | 0.05 | 0.30 | Winter |
| ep_00269 | Chrysophyceae | Clade-I_X | 0.04 | 2 | 1.00 | 1.00 | Summer |

<sup>1</sup> Proportional to the resident microbiota

We identified a total of 18 connector OTUs (featuring relatively low degree and high centrality), which were predominantly associated with summer (5 out of 18) or autumn (6 out of 18), contrasting with hub OTUs, which were associated mostly with winter and spring (**Table 6**). Connectors may be linked to the seasonal transition between main community states (**Figure 1 C & D**) and included several abundant OTUs belonging to *Synechococcus* and SAR11 (**Table 6**). In particular, the SAR11 OTU bp_000007 displayed a relatively high abundance (1.4%), but a degree of 3 (relatively low) and a betweenness centrality of 0.6 (relatively high). In contrast, two protist OTUs displayed low-moderate abundances (ep_00269, Chrysophyceae, abundance 0.04% and en_00161, Syndiniales, abundance 0.4%), low degree <4, but a high betweenness centrality (>0.8; **Table 6**).

## DISCUSSION

Identifying the most important microbes for the functioning of the ocean ecosystem is a challenge, which can be addressed by delineating core microbiotas [4]. Recognizing the most abundant and widespread microbes in the ocean is a step towards knowing the core microbiota. However, this does not take into account the importance that both microbial interactions and microbes with moderate or low abundance may have for the functioning of ecosystems [4, 29, 38]. Considering potential interactions when delineating core microbiotas may not only allow identifying moderate/low abundance taxa that may have important roles in the community but could also allow excluding taxa that are present in several locations but that may not have an important role for community function (e.g., dormant cells or cells being dispersed [8]). Here, we have delineated and analyzed the core microbiota of a coastal ecosystem-based on 10 years of occurrence data considering possible interactions.

To detect the core microbiota, we first identified the resident OTUs, that is, those that occur >30% of the time (i.e. >36 out of 120 months) over a decade. This threshold was selected as it allows for seasonal OTUs that would be present recurrently in at least one season. Analysis of the resident OTU dynamics indicated a clear seasonality (**Figure 1 C-D**), and that the measured environmental factors could explain ∼45% of the resident microbiota variance. The main environmental drivers were temperature and daylength, which is consistent with previous works from the same time-series (BBMO) [34, 39, 40]. These values are lower than what has been reported for bacteria in the English Channel, where daylength explains ∼65% of community variance [17], and higher than what has been reported for entire communities in the time-series SPOT (California, 31%) [41] or SOLA (the Mediterranean Sea, ∼130 km from BBMO; 7-12%) [42]. Daylength may be more important in the English Channel as it has a more pronounced annual variation than at BBMO, whereas the measured differences could reflect a higher coupling of the resident OTUs with environmental variation in BBMO than in SOLA or SPOT. SOLA is characterized by the occasional winter storms that bring nutrients from the sediments to the water column as well as by the freshwater inputs from nearby rivers during flash floods [43], and this could partially explain the differences with BBMO. The importance of daylength and temperature for community dynamics was reflected by niche analyses, which identified two main niches associated with summer and winter at the BBMO, to which ∼50% of the resident OTUs were associated (**Figure 1 E-F)**. Other resident OTUs likely have spring and fall niches as indicated by **Figure 1 C-D**, yet these niches cannot be detected with the used null model analysis, as their preferred temperatures or daylengths will not depart significantly from the randomized mean.

Based on the resident OTUs, we built networks to define the core microbiota. We identified a total of 259 core OTUs (182 bacteria and 77 protists) that represented 64% of the abundance of the resident microbiota and that showed seasonal variation. We could only find supporting evidence from the literature (PIDA database) [21] for 85 associations of the core (6 %), indicating that most of them still need to be validated with direct observation or experimentally. This is not surprising, as the most studied hosts in PIDA are protists from the micro-plankton (>20 µm cell size), which are mostly absent from our pico- and nanoplankton networks. Also, PIDA does not cover Bacteria-Bacteria associations. Nevertheless, the detected core OTUs from BBMO represent a fraction of the core microbiota at this site, since larger microbial size fractions were not sampled. Including these larger size fractions would expand the composition of the core and could unveil additional patterns. For example, in a global ocean network including size fractions >20 µm cell size, protists or small multicellular eukaryotes dominated the interactome [26].

Alpha-/Gammaproteobacteria, Bateroidia, Acidimicrobiia were the main bacterial groups in the core, including also common marine taxa, such as *Synechococcus* or SAR11. The main protists in the core included Syndiniales (parasites), Dinoflagellates, Mammiellales (*Micromonas* and *Bathycoccus*), and diatoms. These taxa are likely the most important in sustaining ecosystem function at BBMO, and probably have similar importance in other coastal areas. Other studies have reported important roles in marine association networks for SAR11 and *Synechococcus* [31, 44]. Syndiniales, Haptophytes, and Dinoflagellates dominated networks in terms of the number of nodes and edges at SPOT, while Mamiellales (*Micromonas* & *Bathycoccus*) and diatoms also had relevant roles [41]. Syndiniales, Dinoflagellates, and Diatoms were also predominant in global ocean networks, which is coherent with our results [26].

Bacteria-Bacteria associations were the most abundant (54%) in the core BBMO microbiota, followed by Bacteria-Protists (31%) and Protist-Protist (11%) associations. Associations tended to occur among bacteria or protists, rather than between them, in the English Channel time-series [17]. However, the study used microscopy to determine protist community composition, while it used 16S-rRNA gene data for analyzing bacteria communities and this might explain the limited number of connections between protists and bacteria. Most associations occurred among protists in a global-ocean network that included a broad range of microbial size-fractions [26]. This suggests that time-series analyses including larger size-fractions may determine a higher proportion of associations among protists, which may turn out to be prevalent.

The core network had “small world” properties (that is, high clustering coefficient and relatively short path lengths) [37] when compared to randomized networks (**Table 3**) or particular subnetworks from size fractions or specific seasons (**Table 5**). The small-world topology is characteristic of many different types of networks [45], including marine microbial temporal or spatial networks [23, 26, 30, 31]. Some of our network statistics were similar to those obtained at SPOT [23, 30], in particular the averages of degree, clustering coefficient, and path length (**Table 3**). Furthermore, the BBMO network had an average path length similar to a global ocean network [26] and also, similarly to this network, the node degree of the BBMO core members was independent of their relative abundances, showing that the associations between core OTUs were not merely a consequence of high prevalence and abundance.

The BBMO core network had a clustering coefficient that was ten times larger than that of an Erdős–Rényi random network of the same size (**Table 3**), which agrees with what was observed at SPOT [23, 30]. The large proportion of positive associations in BBMO networks (∼95%) was in agreement with results from other temporal [23, 41] or large-scale spatial [26] microbiota analyses, where positive associations were also predominant (∼70-98%), although these values include taxa that are not necessarily part of the core. This suggests that interactions such as syntrophy or symbiotic associations are more important than competition in marine microbial systems and that these types of associations may underpin marine ecosystem function. These findings are also coherent with a recent large-scale literature survey that found that ∼47% of the validated associations between protists and bacteria are symbiotic [21]. Nevertheless, it is also possible that common sampling strategies and methodological approaches do not detect a substantial fraction of negative associations. For example, while positive correlations in taxa abundance pointing to positive interactions may be easier to detect, negative associations may be missed due to plummeting species abundances that would prevent establishing significant correlations, or to a delay between the increase and decrease in abundance of interacting taxa that are not synchronized with sampling time. Future studies adapting the sampling scheme to the timing of interactions (e.g., daily or weekly sampling) and the use of other approaches apart from taxa abundances, such as analyses of single-cell genomic data to determine protistan predation, or controlled experiments, will likely generate new insights on negative microbial interactions.

The relatively high clustering coefficient of the core network (compared to a random network) and its short path length indicate that most OTUs are connected through < 3 intermediary OTUs. It has been shown that a large proportion of strong positive associations, as in the BBMO core network, may destabilize communities due to positive feedbacks between species [46]. When a species decreases in abundance as a response to environmental variation, it may pull others with it, generating a cascade effect propagated by the many positive associations in the network. Accordingly, the change of abundance in specific OTUs in one section of the network could affect OTUs in other network sections not necessarily affected directly by the environmental variation. This cascade effect may help to explain a paradox: environmental variables affect the structure of marine microbial communities and consequently association networks. Yet, our and others’ results [17, 18, 23, 26, 30–32] have reported a limited number of associations between environmental variables and network nodes (OTUs). Environmental heterogeneity might affect network structure by acting on a small subset of nodes (OTUs), which would then influence other nodes through cascading interactions facilitated by the highly interconnected nature of the networks as well as positive feedbacks promoted by the high proportion of positive associations [46].

If OTUs susceptible to environmental variation are also highly connected, then their effect on the entire network structure may be larger. In line with this, we found that the connectivity of OTUs associated with environmental variables at BBMO (49 OTUs out of 259) had a mean degree of ∼25 (SD ∼14), while for all the 259 OTUs of the core network, the mean degree was ∼11 (SD ∼13). The seasonal dynamics of the BBMO microbiota may partially be driven by a subset of OTUs that vary with environmental factors (e.g. temperature, daylength). These may exert a destabilizing influence over the entire community over time, promoting the annual turnover of communities and networks.

Most core OTUs (98%) showed a clear preference for one season. Interestingly, the distribution of core OTUs among the seasons was uneven, with 61% of these OTUs showing a winter preference. Network connectivity at BBMO was correspondingly heterogeneous between seasons, peaking in winter and remaining low in the other seasons. Specifically, the winter subnetwork included ∼92% of the seasonal edges. This indicates that winter associations are not only specific (i.e. they do not tend to change partners), but they also have a relatively high recurrence (otherwise, winter networks would be smaller). A higher similarity between winter communities when compared to other seasons was also indicated by our ordination analyses of the resident OTUs (**Figure 1**), as well as by studies of the entire protist community at BBMO [34] or whole community analyses at SPOT [23].

The structure of communities is determined by the interplay of selection, dispersal, speciation, and ecological drift [47]. Our results indicate that selection, a deterministic process, is stronger in winter, leading to winter sub-communities that tend to be more similar between each other than to communities from other seasons. Given that we have removed edges associated with the measured environmental variables, we do not expect that the identified edges between winter OTUs represent selection associated to these variables (e.g. low temperature). Consequently, winter edges may represent associations linked to unmeasured variables or ecological interactions that may be more likely to develop during winter due to stronger environmental selection. Due to weaker selection in other seasons species occurrence would display less recurrent (or more random) patterns, preventing specific associations to be formed. This also suggests that ecological redundancy changes over time, and is lower in winter compared to the other seasons (even though the number of OTUs is larger in winter). A reduction in redundancy may also promote strong ecological interactions in winter.

The existence of subsets of species that interact more often between themselves than with other species (modules), is characteristic of biological networks, and can contribute to overall network stability [48, 49]. Modules can represent divergent selection, niches, the clustering of evolutionary closely related species or co-evolutionary units [50, 51]. Modules in the core BBMO network (total 12) included positive associations between diverse taxa, and could represent divergent selection, driven by unmeasured environmental variables, or examples of syntrophic or symbiotic interactions between microbes from different taxonomic groups.

Most BBMO modules included diverse lifestyles (heterotrophs, mixotrophs, phototrophs, parasites), similar to what has been observed at SPOT [41]. Yet, a number of modules appeared to be predominantly heterotrophic or autotrophic (**Table S8, Additional file 1**). Some modules included OTUs from the same species, such as Module 4 in the picoplankton, which included several SAR11 Clade I OTUs, and Module 7 of the nanoplankton, which included several *Synechococcus* OTUs. These modules could reflect similar niches, associated with unmeasured variables, or the dependence on metabolites produced by other organisms (auxotrophy). There is evidence of auxotrophy for both SAR11 (e.g. thiamin, glycine)[52–54] and *Synechococcus* (e.g. cobalamin) [55]. Recently it has been observed in co-culture experiments that *Prochlorococcus* may fulfill some metabolic requirements of SAR11, promoting the growth of the latter in a commensal relationship [56]. In our analyses of the BBMO core microbiota, we did not find strong associations between SAR11 and *Prochlorococcus* or the more abundant relative, *Synechococcus*. Yet, SAR11 formed strong associations with a plethora of taxa with which could potentially have commensal relationships.

The overall importance of the observed modules was indicated by the total abundance of their constituent OTUs (24% of the reads compared to the resident microbiota). Most of the modules at BBMO were associated with a single season, suggesting that they reflect seasonal niches. Since these modules were inferred over 10 years, they represent recurrent network features. Chafee et al. [57] also identified season-specific modules in a 2-year time series in the North Sea (Helgoland), including samples taken weekly or bi-weekly. These modules were much larger than ours, and they may also include environmentally-driven edges. Nevertheless, the Helgoland modules seem to be driven by eutrophic (spring & summer) vs. oligotrophic (autumn & winter) conditions in this location. In contrast, the BBMO modules, displayed weaker correlations with nutrients and seem to be influenced by temperature and daylength (**Figure 5**). Differences in the sampling scheme between Helgoland and BBMO ((bi)weekly vs. monthly) as well as between both locations (different seas and latitudes, affecting temperature and daylength) may explain these differences.

Keystone species have a high influence in ecosystems relative to their abundance [58]. Network analyses may help to identify them [24, 59], yet, there is no clear consensus of what network features are the best unequivocal indicator of keystone species [60–62]. Therefore, we focused on identifying central OTUs (hubs or connectors) that may be important for ecosystem function [22, 24] and could represent keystone species. We identified 13 hubs in the BBMO core network with moderate-low abundances (<1%) and high degree (26-60) that were associated with winter or spring. These moderate-low abundance OTUs may affect nutrient cycling directly [63] or indirectly, by affecting other OTUs with higher abundance. The putative stronger selection exerted by low temperatures and short daylengths during winter and early spring, as compared to summer and autumn, may lead to a higher species recurrence [34], larger networks, and possibly, more hubs. An OTU of the abundant picoalgae *Bathycoccus* (en_00092) was identified as a winter hub, which is consistent with reported *Bathycoccus* abundance peaks in late winter (February-March) in both BBMO [64] and the nearby station SOLA [42]. This *Bathycoccus* hub may be associated with diverse taxa, such as prokaryotes that may benefit from algal exudates [65] or even via mixotrophy [66]. In agreement with this, out of the 42 associations of this hub OTU, 25 were with bacteria and the rest with protists.

In contrast to hubs, connector OTUs were predominantly associated with warmer waters, that is, summer and autumn, and may represent transitions in community states. This was consistent with the associations observed in an abundant *Synechococcus* connector OTU (bp_000001, **Table 6**). This OTU was predominant in summer-autumn, in agreement with previous BBMO reports [36, 67], but it was associated with other OTUs from spring (negative association with bp_000017), winter (negative association with bp_000039), summer (positive association with bp_000087, bp_000012) and autumn (positive association with bp_000022), thus likely holding a central position in the network. Another abundant spring connector OTU (SAR11 Clade Ia, bp_000002), featured only two connections to spring (positive association with bp_000007) and summer (positive association with bp_000046) OTUs.

## CONCLUSION

Our decade-long analysis of the dynamics of a microbiota populating a time-series in the Mediterranean Sea allowed us to determine the interconnected core microbiota, which likely includes several microbes that are important for the functioning of this coastal ecosystem. We found a relatively small core microbiota that displayed seasonal variation, with a heterogeneous distribution of associations over different seasons, indicating different degrees of recurrence and selection strength over the year. Future analyses of other core marine microbiotas will determine how universal are the patterns found in BBMO. These studies will be crucial to determine potential long-term effects of climate change on the architecture of the interaction networks that underpin the functioning of the ocean ecosystem.

## METHODS

### Study site and sampling

Surface water (∼1 m depth) was sampled monthly from January 2004 to December 2013 at the Blanes Bay Microbial Observatory (BBMO) in the Northwestern Mediterranean Sea (41°40’N, 2°48’E) [**Figure 1A**]. The BBMO is an oligotrophic coastal site ∼1 km offshore with ∼20 m depth and with limited riverine or human influence [36]. Seawater was pre-filtered with a 200 µm nylon mesh and then transported to the laboratory in 20 L plastic carboys and processed within 2 hours. Microbial plankton from about 6 L of the pre-filtered seawater was separated into two size fractions: picoplankton (0.2-3 µm) and nanoplankton fraction (3-20 µm). To achieve this, the seawater was first filtered through a 20 µm nylon mesh using a peristaltic pump. Then the nanoplankton (3-20 µm) was captured on a 3 µm pore-size polycarbonate filter. Subsequently, a 0.2 µm pore-size Sterivex unit (Millipore, Durapore) was used to capture the picoplankton (0.2-3 µm). Sterivex units and 3 µm filters were stored at −80 °C until further processed. The sequential filtering process aimed to capture free-living bacteria and picoeukaryotes in the 0.2-3 µm size fraction (picoplankton), and particle/protist-attached bacteria or nanoeukaryotes in the 3-20 µm fraction (nanoplankton). The 3µm filter was replaced if clogging was detected; DNA from all 3µm filters from the same sample were extracted together.

A total of 15 contextual abiotic and biotic variables were considered for each sampling point: Daylength (hours of light), Temperature (°C), Turbidity (estimated as Secchi disk depth [m]), Salinity, Total Chlorophyll a [Chla] (μg/l), PO_4_^3-^ (μM), NH_4_^+^ (μM), NO_2_^-^ (μM), NO_3_^-^ (μM), SiO_2_ (μM), abundances of Heterotrophic prokaryotes [HP] (cells/ml), *Synechococcus* (cells/ml), Total photosynthetic nanoflagellates [PNF; 2-5µm size] (cells/ml), small PNF (2µm; cells/ml) and, Heterotrophic nanoflagellates [HNF] (cells/ml) [**Figure 1B**]. Water temperature and salinity were sampled *in situ* with a SAIV A/S SD204 CTD. Inorganic nutrients (NO_3_^-^, NO_2_^-^, NH_4_^+^, PO_4_^3-^, SiO_2_) were measured using an Alliance Evolution II autoanalyzer [68]. Cell counts were done by flow cytometry (heterotrophic prokaryotes, *Synechococcus*) or epifluorescence microscopy (PNF, small PNF and HNF). See Gasol *et al.* [36] for specific details on how other variables were measured. Environmental variables were z-score standardized before running statistical analysis.

### DNA extraction, sequencing, and metabarcoding

DNA was extracted from the filters using a standard phenol-chloroform protocol [69], purified in Amicon Units (Millipore), and quantified and qualitatively checked with a NanoDrop 1000 Spectrophotometer (Thermo Fisher Scientific). Eukaryotic PCR amplicons were generated for the V4 region of the 18S rDNA (∼380 bp), using the primer pair TAReukFWD1 and TAReukREV3 [70]. The primers Bakt_341F [71] and Bakt_806RB [72] were used to amplify the V4 region of the 16S rDNA. PCR amplification and amplicon sequencing were carried out at the Research and Testing Laboratory (http://rtlgenomics.com/) on the *Illumina* MiSeq platform (2×250 bp paired-end sequencing). DNA sequences and metadata are publicly available at the European Nucleotide Archive (http://www.ebi.ac.uk/ena; accession numbers PRJEB23788 for 18S rRNA genes & PRJEB38773 for 16S rRNA genes).

A total of 29,952,108 and 16,940,406 paired-end *Illumina* reads were produced for microbial eukaryotes and prokaryotes respectively. Adapters and primers were removed with Cutadapt v1.16 [73]. DADA2 v1.10.1 [74] was used for quality control, trimming, and inference of Operational Taxonomic Units (OTUs) as Amplicon Sequence Variants (ASVs). For both microbial eukaryotes and prokaryotes, the Maximum number of expected errors (MaxEE) was set to 2 and 4 for the forward and reverse reads respectively. No ambiguous bases (Ns) were allowed. Microbial eukaryotic sequences were trimmed to 220 bp (forward) and 190 bp (reverse), while prokaryotic sequences were trimmed to 225 bp (both forward and reverse reads). A total of 28,876 and 19,604 OTUs were inferred for microbial eukaryotes and prokaryotes respectively.

OTUs were assigned taxonomy using the naïve Bayesian classifier method [75] together with the SILVA version 132 [76] database as implemented in DADA2. Eukaryotic OTUs were also BLASTed [77] against the Protist Ribosomal Reference database (PR^2^, version 4.10.0; [78]). When the taxonomic assignments for the eukaryotes disagreed between SILVA and PR^2^, the conflict was resolved manually by inspecting a pairwise alignment of the OTU and the closest hits from the two databases. OTUs assigned to Metazoa, Streptophyta, nucleomorph, chloroplast, and mitochondria were removed before further analysis. Archaea were removed from downstream analyses as the used primers are not optimal for recovering this domain [79].

Each sample (corresponding to a specific gene, size fraction, and timepoint) was subsampled with the *rrarefy* function from the R package *Vegan* [80] to 4,907 reads, corresponding to the number of reads in the sample with the lowest sequencing depth, to normalize for different sequencing depth between samples. OTUs present in <10% of the samples were removed. After quality control and rarefaction, the number of OTUs was 2,926 (1,561 bacteria, and 1,365 microeukaryotes; **Table 1**).

Due to a suboptimal sequencing of the amplicons, we did not use nanoplankton samples of bacteria and protists from the period May 2010 to July 2012 (27 samples) as well as March 2004 and February 2005. OTU read abundance for samples with missing values were estimated using seasonally aware missing value imputation by weighted moving average for time series as implemented in the R package *imputeTS* [81].

Cell/particle dislodging or filter clogging during the sequential filtration process may affect the taxonomic diversity observed in the different size fractions, with nanoplankton DNA leaking into the picoplankton fraction, or picoplankton DNA getting stuck in the nanoplankton fraction. To minimize the effects of cell/particle dislodging or filter clogging on the diversity recovered from the different size fractions, we calculated the sequence-abundance ratio for OTUs appearing in both pico- and nano-plankton fractions. When the ratio exceeded 2:1, we removed the OTU from the size fraction with the lowest number of reads. After subsampling and filtering the OTU tables were joined for each time point, and since the samples had been normalized to the same sequencing depth, we calculated the relative read abundance for the OTUs for each year and aggregated over the corresponding months along the 10 years for the resident microbiota. This means that the relative abundance for both domains and size fractions sums up to 1 for each month across ten years.

### Resident microbiota

We defined *ad hoc* the resident microbiota as the set of OTUs present in >30% of the samples over 10 years (that is, present in >36 months, not necessarily consecutive). This value was chosen as it allows for seasonal OTUs, which may only be present 3-4 months each year, and still be considered as part of the resident microbiota. The residents included 355 eukaryotic and 354 bacteria OTUs (**Table 1**), and excluded a substantial amount of rare OTUs, which can cause spurious correlations during network construction due to sparsity [i.e. too many zeros] [22]. The relative abundance of the taxonomic groups included in the resident microbiota was fairly stable from year to year (**Figure 3**).

### Environmental variation and resident OTUs

All possible correlations among the measured environmental variables and resident OTU richness and abundance were computed in R and plotted with the package *corrplot*. Only significant Pearson correlation coefficients were considered (p<0.01), and the p-values were corrected for multiple inference (Holm’s method) using the function *rcorr.adjust* from the R package *RcmdrMisc*. Unconstrained ordination analyses were carried out using NMDS based on Bray Curtis dissimilarities between samples including resident OTUs only. Environmental variables were fitted to the NMDS using the function *envfit* from the R package *Vegan* [80]. Only variables displaying a significant correlation (p<0.05) were considered. Constrained ordination was performed using distance-based redundancy analyses (dbRDA) in *Vegan*, considering Bray Curtis dissimilarities between samples including resident OTUs only. The most relevant variables for constrained ordination were selected by stepwise model selection using 200 permutations, as implemented in *ordistep* (*Vegan*). Ordinations were plotted using the R package *ggplot2* and *ggord*. The amount of community variance explained by the different environmental variables was calculated with *Adonis* (Vegan) using 999 permutations. Resident OTUs displaying niche preference in terms of Temperature and Daylength, the most important environmental variables, were determined using the function *niche.val* from the R package *EcolUtils* with 1,000 permutations.

### Delineation of seasons

Seasons were defined following Gasol *et al.* [36] with a small modification: months with water temperature (at the sampling time) >17 °C and daylength >14 h d^−1^ were considered to be summer. Months with water temperature <17 °C and < 11 h d^−1^ of daylength were considered to be winter. Months with water temperature >17°C and daylength <14 h d^−1^ were considered as autumn, while months with water temperature <17°C and > 11 h d^−1^ of daylength were considered to be spring. The indicator value [82] was calculated using the R package *labdsv* [83] to infer OTU seasonal preference.

### Core microbiota delineated using networks

The OTU table together with the 15 environmental variables were used to construct association networks using extended Local Similarity Analysis (eLSA) [84–86]. eLSA was run on the OTU table with subsampled reads with default normalization: a z-score transformation using the median and median absolute deviation. P-value estimations were run under a mixed model that performs a random permutation test of a co-occurrence only if the theoretical p-values for the comparison are <0.05. Bonferroni false discovery rate (q) was calculated for all edges based on the p-values using the *p.adjust* package in R.

To detect environmentally-driven associations between OTUs induced by the measured environmental variables we used the program EnDED [87]. Environmentally-driven associations indicate similar or different environmental preferences between OTUs and not ecological interactions. In short, EnDED evaluates associations between two OTUs that are both connected to the same environmental variable based on a combination of four methods: *Sign Pattern*, *Overlap, Interaction Information,* and *Data Processing Inequality.* These methods use the sign (positive or negative) and the duration of the association, the relative abundance of OTUs as well as environmental parameters to determine if an association is environmentally-driven. If the four methods agreed that an association was environmentally-driven, then it was removed from the network. The initial number of edges was 199,937, of which 180,345 were OTU-OTU edges that were at least in one triplet with an environmental parameter. In total 65,280 (∼33%) edges in the network were identified as indirect by EnDED and removed. Afterward, only edges representing the strongest associations (i.e., absolute local similarity score |LS| > 0.7, Spearman correlation |ρ| > 0.7, P<0.001 and Q<0.001) and nodes representing the resident OTUs were retained for downstream analysis and are hereafter referred to as “core associations”. Those OTUs participating in core associations were defined as core OTUs, although their involvement in ecological interactions need further experimental validation. Both core associations and core OTUs constitute the “core network”, which also represents the core microbiota (both “core network” and “core microbiota” are used indistinctively). The core network was randomized using the Erdős–Rényi model [88], using 262 nodes and 1,411 edges.

For the core network, we calculated: 1) *Density:* quantifies the proportion of actual network connections out of the total number of possible connections, 2) *Transitivity* or *Clustering coefficient:* measures the probability that nodes connected to a node are also connected, forming tight clusters, 3) *Average path length:* mean number of steps (edges) along the shortest paths for all possible pairs of nodes in the network (a low average path length indicates that most species in the network are connected through a few intermediate species), 4) *Degree:* number of associations per node, 5) *Betweenness centrality:* measures how often an OTU (node) appears on the shortest paths between other OTUs in the network, 6) *Closeness centrality:* indicates how close a node is to all other nodes in a network, 7) *Cliques:* refers to sets of interconnected nodes where all possible connections are realized, 8) *Modularity:* measures the division of a given network into modules (that is, groups of OTUs that are highly interconnected between themselves).

The Degree, Betweenness centrality and Closeness centrality were used to identify central OTUs using *ad hoc* definitions. “Hub” OTUs were those with a score above the average for the three statistics and were normally among the top 25% in each score [22, 62, 89]. Specifically, hub OTUs featured a degree >24, Betweenness centrality >0.03 and Closeness centrality >0.3. Similarly, “connector” OTUs were defined as those featuring a relatively low degree and high centrality and could be seen as elements that connect different regions of a network or modules [50]. Connector OTUs featured a degree <5, Betweenness centrality > 0.03 and Closeness centrality >0.2. Network statistics were calculated with *igraph* in R [90], Gephi [91] and Cytoscape v3.6.1 [92]. Visualizations were made in Cytoscape v3.6.1. Modules in the core network were identified with MCODE [93].

## DECLARATIONS

### Ethics approval and consent to participate

Not applicable

### Consent for publication

Not applicable

### Availability of data and materials

DNA sequences and metadata are publicly available at the European Nucleotide Archive (http://www.ebi.ac.uk/ena; accession numbers PRJEB23788 [18S rRNA genes] & PRJEB38773 [16S rRNA genes]).

### Authors’ contributions

AKK & RL designed the study. JMG, RM organized sampling. VB, CRG & IF collected samples, extracted the DNA, and organized its sequencing. AKK, RL & ID analyzed the data, while JMG, RM, IF, CRG & EG, provided contextual ecological or environmental pre-processed data. AKK, MFMB & RL interpreted the results. AKK & RL wrote the manuscript. All authors contributed substantially to manuscript revisions. All authors read and approved the final manuscript.

## Acknowledgements

We thank all members of the Blanes Bay Microbial Observatory sampling and analyses team. Bioinformatics analyses were performed at the MARBITS platform of the Institut de Ciències del Mar (ICM; http://marbits.icm.csic.es). We thank the CSIC Open Access Publication Support Initiative through the Unit of Information Resources for Research (URICI) for helping to cover publication fees.

## ADDITIONAL FILES

**Additional file 1: Table S1**

Relative abundance of bacterial and protistan lineages that are part of the resident and core microbiotas.

**Additional file 1: Table S2**

Relative abundance of core bacterial taxa.

**Additional file 1: Table S3**

Relative abundance of core eukaryotic taxa.

**Additional file 1: Table S4**

Indicator value for core OTUs in the picoplankton. Sorted by season/kingdom and relative amplicon abundance.

**Additional file 1: Table S5**

Indicator value for core OTUs in the nanoplankton. Sorted by season/ kingdom and relative amplicon abundance.

**Additional file 1: Table S6**

Core OTUs without seasonal preference.

**Additional file 1: Table S7**

Module description.

**Additional file 1: Table S8**

OTUs within modules.

**Additional file 2: Figure S1**

**Panel A** shows the full network constructed with the resident microbiota (that is, OTUs present in >30% of the samples over 10 years; Table 1). **Panel B** displays network elements that were removed as they did not fulfill the cut-offs (that is, highly significant correlations (P & Q <0.001), local similarity scores >|0.7| and Spearman correlations >|0.7|).

**Additional file 3: Figure S2**

OTU relative abundance vs. degree shows no relationship in the core network.

## Supplementary Figures

**Figure S1.**
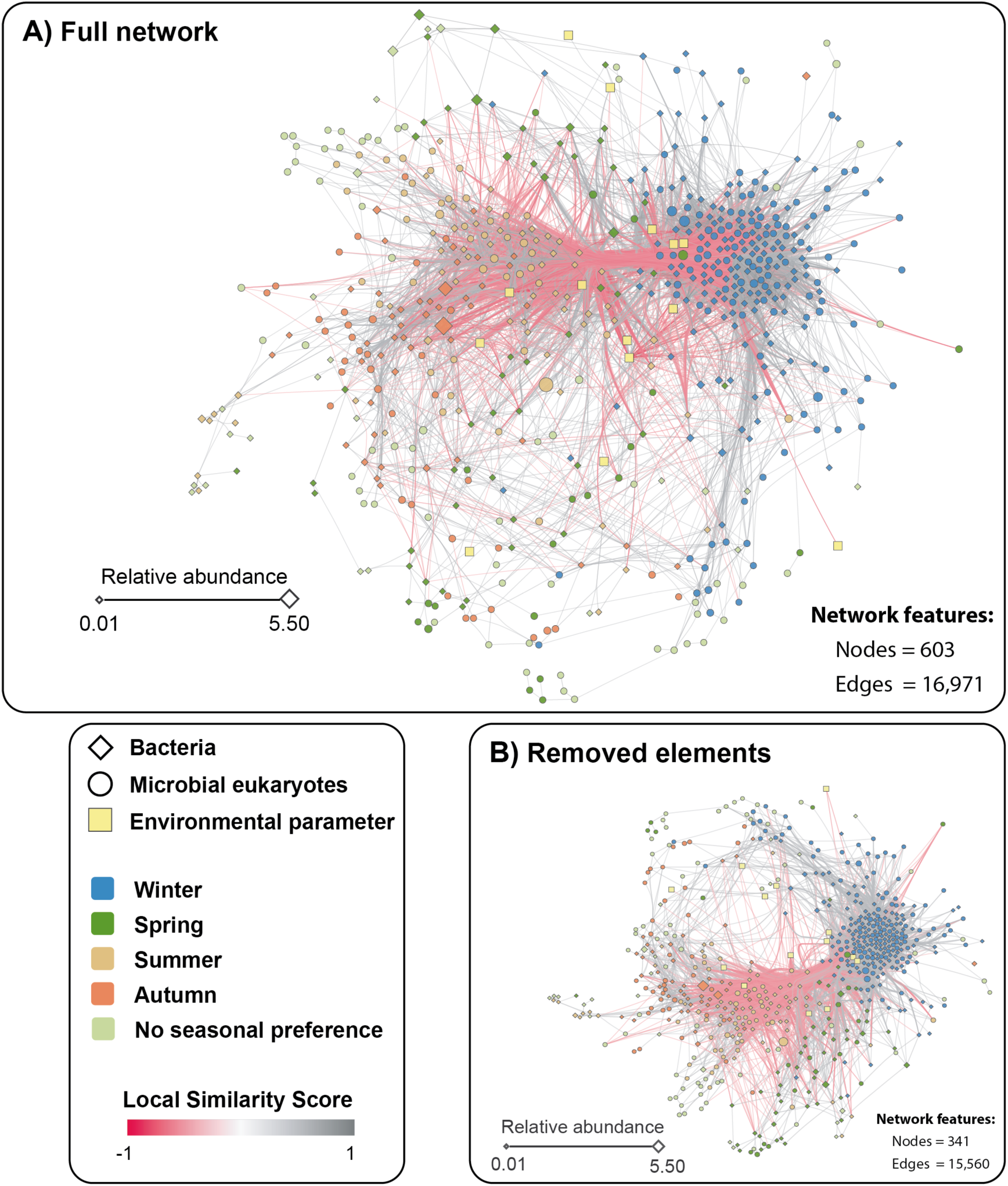
**Panel A** shows the full network constructed with the resident microbiota (that is, OTUs present in >30% of the samples over 10 years; Table 1). **Panel B** displays network elements that were removed as they did not fulfill the cut-offs (that is, highly significant correlations (P & Q <0.001), local similarity scores >|0.7| and Spearman correlations >|0.7|).

**Figure S2.**
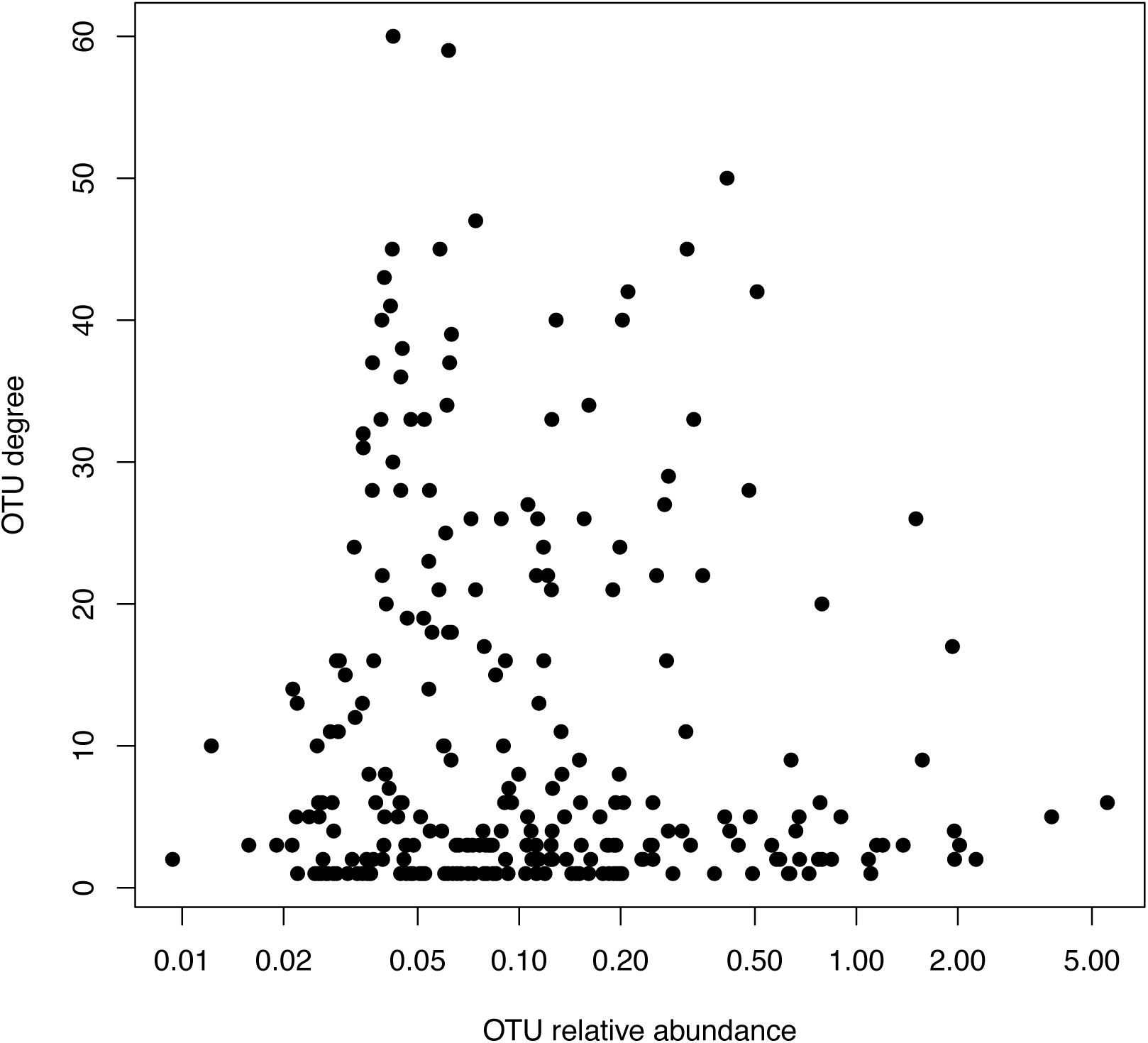
OTU relative abundance vs. degree shows no relationship in the core network.

